# Neutrophil Profiles of Pediatric COVID-19 and Multisystem Inflammatory Syndrome in Children

**DOI:** 10.1101/2021.12.18.473308

**Authors:** Brittany P. Boribong, Thomas J. LaSalle, Yannic C. Bartsch, Felix Ellett, Maggie E. Loiselle, Jameson P. Davis, Anna L. K. Gonye, Soroush Hajizadeh, Johannes Kreuzer, Shiv Pillai, Wilhelm Haas, Andrea Edlow, Alessio Fasano, Galit Alter, Daniel Irimia, Moshe Sade-Feldman, Lael M. Yonker

**Affiliations:** Mucosal Immunology and Biology Research Center, Massachusetts General Hospital; Boston, USA; Department of Pediatrics, Massachusetts General Hospital; Boston, USA; Department of Medicine, Harvard Medical School; Boston, USA; Center for Cancer Research, Department of Medicine, Massachusetts General Hospital; Boston, USA; Broad Institute of MIT and Harvard; Cambridge, USA; Ragon Institute of MGH, MIT and Harvard; Cambridge, USA; BioMEMS Resource Center, Department of Surgery, Massachusetts General Hospital, Shriners Burns Hospital, Harvard Medical School; Boston, USA; Department of Obstetrics and Gynecology, Division of Maternal-Fetal Medicine; Boston, USA; Vincent Center for Reproductive Biology, Massachusetts General Hospital; Boston, USA; European Biomedical Research Institute of Salerno (EBRIS); Salerno, Italy

## Abstract

Multisystem Inflammatory Syndrome in Children (MIS-C) is a delayed-onset, COVID-19-related hyperinflammatory systemic illness characterized by SARS-CoV-2 antigenemia, cytokine storm and immune dysregulation; however, the role of the neutrophil has yet to be defined. In adults with severe COVID-19, neutrophil activation has been shown to be central to overactive inflammatory responses and complications. Thus, we sought to define neutrophil activation in children with MIS-C and acute COVID-19. We collected samples from 141 children: 31 cases of MIS-C, 43 cases of acute pediatric COVID-19, and 67 pediatric controls. We found that MIS-C neutrophils display a granulocytic myeloid-derived suppressor cell (G-MDSC) signature with highly altered metabolism, which is markedly different than the neutrophil interferon-stimulated gene (ISG) response observed in pediatric patients during acute SARS-CoV-2 infection. Moreover, we identified signatures of neutrophil activation and degranulation with high levels of spontaneous neutrophil extracellular trap (NET) formation in neutrophils isolated from fresh whole blood of MIS-C patients. Mechanistically, we determined that SARS-CoV-2 immune complexes are sufficient to trigger NETosis. Overall, our findings suggest that the hyperinflammatory presentation of MIS-C could be mechanistically linked to persistent SARS-CoV-2 antigenemia through uncontrolled neutrophil activation and NET release in the vasculature.

**One Sentence Summary:** Circulating SARS-CoV-2 antigen:antibody immune complexes in Multisystem Inflammatory Syndrome in Children (MIS-C) drive hyperinflammatory and coagulopathic neutrophil extracellular trap (NET) formation and neutrophil activation pathways, providing insight into disease pathology and establishing a divergence from neutrophil signaling seen in acute pediatric COVID-19.

## INTRODUCTION

Neutrophil activation is a central component of the immune response against SARS-CoV-2 infection; however, excessive neutrophil hyperactivation can contribute to severe COVID-19 in adults (*1–4*). The role of neutrophils in pediatric SARS-CoV-2 infections has not been defined. Children are less likely to suffer serious symptoms of acute COVID-19, but some children can develop a severe, life-threatening hyperinflammatory illness called Multisystem Inflammatory Syndrome in Children (MIS-C) weeks to months after the resolution of upper respiratory tract symptoms (*5*). MIS-C is defined by sepsis-like physiology with high fevers, systemic inflammation, and multi-organ involvement, likely caused by SARS-CoV-2 antigenemia originating from gastrointestinal sources of virus leaking across a permeable mucosal barrier into circulation (*6*). Children with MIS-C display signs of vascular injury, including elevated C-reactive protein, erythrocyte sedimentation rate, and D-dimer. Eighty percent of children with MIS-C can develop cardiac complications including myocarditis, ventricular dysfunction, and coronary aneurysms (*7, 8*), although mechanisms of cardiovascular injury are unclear.

Neutrophils play a major role in innate immune responses but hyperactivation of neutrophils has been shown to be detrimental in severe diseases including sepsis, ARDS, and severe acute COVID-19 in adults (*1, 9, 10*). Not only can hyperactivation of polymorphonuclear leukocytes (PMNs) result in vascular and organ injury, but they can also contribute to hypercoagulation and thrombus formation (*11*). Understanding neutrophil activation in acute pediatric COVID-19 and MIS-C is essential to developing clinical guidelines and advances in treatment recommendations.

Extensive immune profiling has shown distinct hyperinflammatory patterns distinguishing acute pediatric COVID-19 from MIS-C (*12–14*); however, very little information on neutrophil activation has been included in these reports. This paucity of research on neutrophil activation in acute pediatric COVID-19 and MIS-C may be due to technical challenges with studying neutrophils: neutrophils have a very short lifespan and are extensively damaged by freezing. In contrast with peripheral blood mononuclear cells (PBMCs), neutrophils require isolation and fixation or analysis within hours of blood collection. Thus, there is still a need for a thorough profiling of neutrophils in pediatric COVID-19 and MIS-C to elucidate their role in disease pathogenesis.

Here, we isolated neutrophils immediately after blood collection from children with acute COVID-19, MIS-C and healthy controls and extensively profiled gene expression, protein production, and neutrophil functionality to define neutrophil responses driving these distinct SARS-CoV-2 disease states in children.

## RESULTS

### Cohort characteristics

Blood was collected from 141 children, including 43 children with COVID-19 confirmed by RT-PCR **(Table S1)**, 31 children clinically diagnosed with MIS-C **(Table S2)**, and 67 non-COVID-19 pediatric controls. Subsets of each group were used for bulk RNA sequencing (RNA-seq) (*n* = 48), proteomics (*n* = 55), cell-free DNA measurements (*n* = 48), and neutrophil extracellular traps (NET) assays (*n* = 14) (**Fig. 1**; **Table 1**). The average age of the acute pediatric COVID-19 cohort, MIS-C cohort, and non-COVID-19 pediatric control cohort was 14.2 years, 7.4 years, and 10.2 years, respectively. Sex, race, and ethnicity are shown in **Table 1**.

**Table 1.**
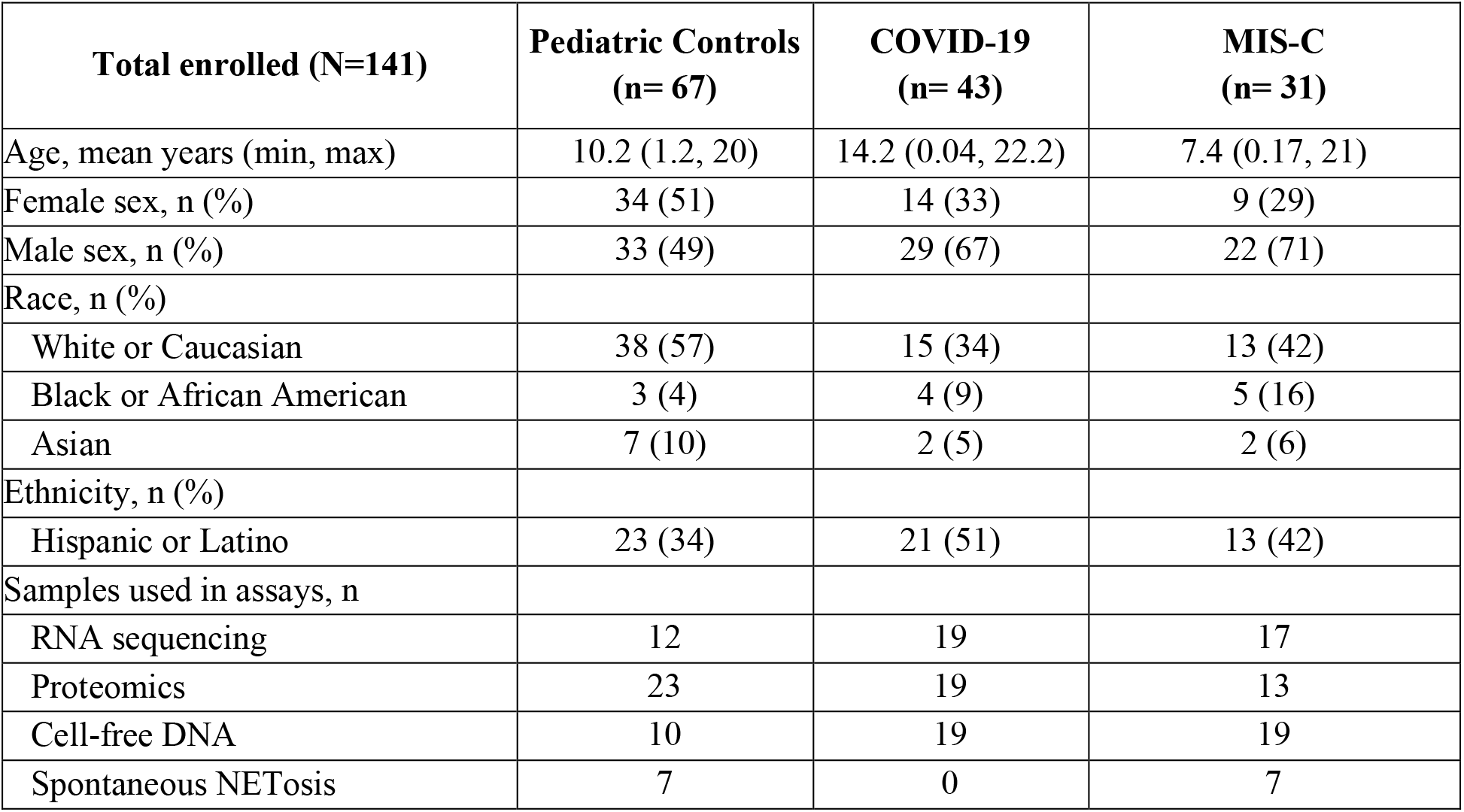
Patient demographics and distribution of pediatric blood samples used for each assay.

**Fig. 1.**
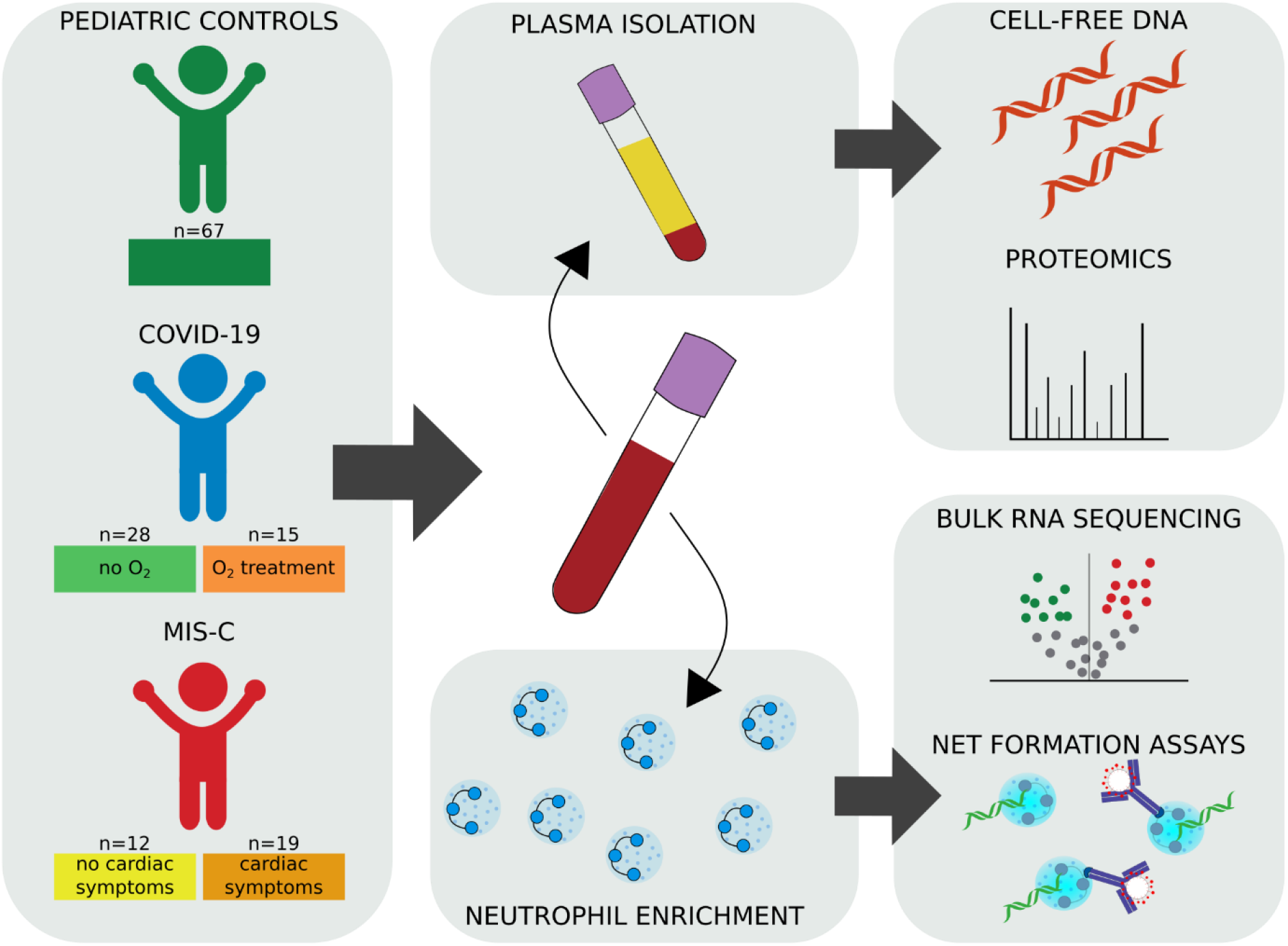
Overview of cohort and experimental schematic. Whole blood samples were collected from three cohorts: (1) pediatric controls (*n* = 67) (2) pediatric acute COVID-19 infection (*n* = 43) and (3) multisystem inflammatory syndrome in children (MIS-C) (*n* = 31). Plasma collected from whole blood samples were used in proteomic assays to quantify levels of protein and cfDNA. Isolated neutrophils were used in bulk RNA-sequencing and NETosis assays. Patient samples were used in individual or multiple assays.

### Transcriptional profiling of neutrophils from pediatric COVID-19 and MIS-C

To confirm successful negative selection enrichment of neutrophils, we used a digital cytometry method, CIBERSORTx, to estimate the fractions of major cell type lineages in each bulk RNA-seq sample (*15*). We used the single-cell RNA-sequencing COVID-19 Bonn Cohort 2 reference dataset to generate cell-type-specific signatures for mature neutrophils, immature neutrophils, monocytes, T/NK cells, B cells, and plasmablasts (*2*). The total neutrophil fraction was defined as the sum of mature and immature neutrophils. High purity of neutrophils was confirmed in all samples that passed quality control (**Fig. S1A**, **Data File S1**), and the majority of samples had high mature-to-immature neutrophil ratios. Notably, there were no significant differences in any CIBERSORTx fractions between any of the disease categories (**Fig. S1B**). Additionally, flow cytometry confirmed high neutrophil purity with minimal monocyte and lymphocyte contamination (**Fig. S1C**).

Uniform manifold approximation and projection (UMAP) visualization of bulk RNA-seq samples did not reveal distinct groupings based on disease status; rather, the landscape was defined by a larger mature neutrophil grouping and a smaller immature neutrophil group (**Fig. S1D-E**).

### Acute COVID-19 infection is associated with neutrophil interferon response signatures in children

To identify differentially expressed genes and pathways in neutrophils associated with acute SARS-CoV-2 infection in children, we performed differential expression analysis and gene set enrichment analysis (GSEA) between COVID-19-positive samples and healthy samples (**Data File S2**). The most highly upregulated gene in COVID-19-positive samples was *CCRL2*, a gene critical for CXCR2-dependent neutrophil recruitment and β2-integrin activation (*16*) (**Fig. 2A**). Following closely were other genes implicated in neutrophil chemotaxis, activation, and type I and II interferon response such as *ELMO2*, *GPR84*, *IRF7*, *IFIT3*, and *MX1*. Overall, COVID-19-positive samples showed strong enrichment of pathways involved in viral response, including response to IFNγ and IFNα (**Fig. 2B, Data File S2**).

**Fig. 2.**
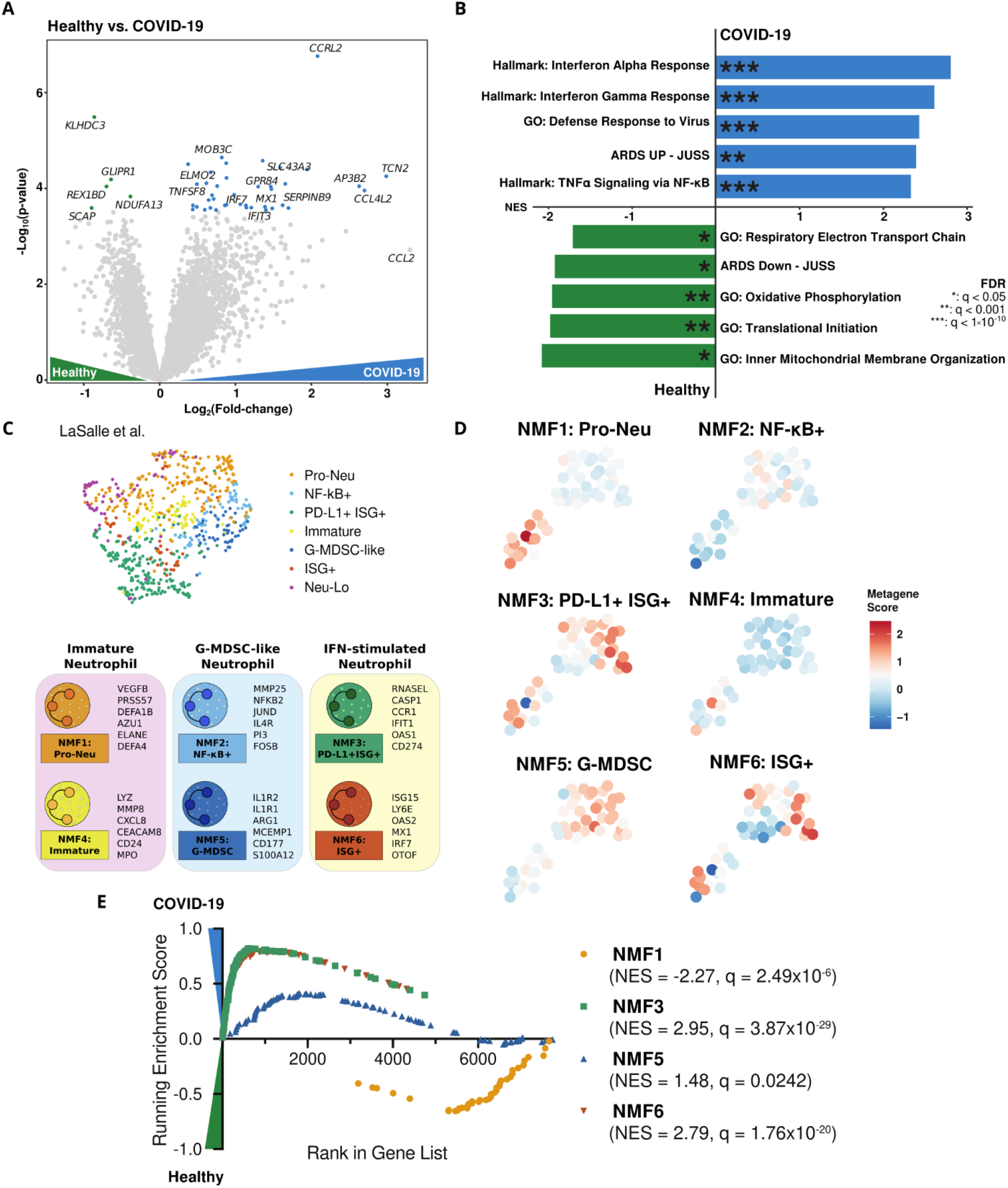
Acute pediatric COVID-19 neutrophils are marked by a robust interferon-stimulated gene signature. (**A)** Volcano plot showing differentially expressed genes between pediatric acute COVID-19 and healthy control samples. Color-coded points indicate genes that pass FDR correction with q < 0.05. **(B)** Gene set enrichment analysis for the differentially expressed genes in **(A)**. Bar lengths correspond to normalized enrichment score (NES), with positive NES values indicating enrichment in COVID-19 and negative values indicating enrichment in pediatric healthy controls. Asterisks denote significance as defined in the figure. **(C)** UMAP of neutrophil bulk RNA-seq samples from adults with acute COVID-19 and healthy controls from LaSalle et al (*1*). UMAP is color-coded by cluster designation. Below, schematic illustrating several top genes associated with each neutrophil subtype. Neu-Lo; samples excluded from clustering based on estimated low neutrophil fraction by CIBERSORTx. **(D)** UMAPs of all neutrophil samples that were sequenced with bulk RNA-seq. Samples are colored according to their expression score of each neutrophil state metagene. **e**. GSEA enrichment plots for the NMF1 (Pro-Neu), NMF3 (PD-L1+ ISG+), NMF5 (G-MDSC), and NMF6 (ISG+) signatures, the four NMF neutrophil signatures which passed FDR correction.

In order to identify bulk neutrophil gene expression subtypes involved in response to SARS-CoV-2 infection in children, we utilized the bulk neutrophil gene signatures defined by an identical protocol in a large adult cohort, as well as previously defined neutrophil signatures from single-cell and bulk RNA-seq data (*10, 17*). Pre-defined signatures included progenitor neutrophils (NMF1: Pro-Neu), NF-κB-activated neutrophils (NMF2: NF-κB+), interferon-stimulated neutrophils with high expression of PD-L1 (NMF3: PD-L1+ ISG+), immature activated neutrophils (NMF4: Immature), granulocytic myeloid-derived suppressor cell-like (G-MDSC) neutrophils (NMF5: G-MDSC), and a second interferon-stimulated neutrophil signature with lower expression of PD-L1 and a non-overlapping set of interferon-stimulated genes (ISGs) (NMF6: ISG+) (*1, 10, 17*) (**Fig. 2C**). When scoring each sample, we found that many of the NMF signatures produced groupings on the UMAP (**Fig. 2D**), indicating the utility of these bulk signatures in defining neutrophil state signatures in children with COVID-19.

Next, we performed gene set enrichment analysis comparing neutrophil subtype signature gene sets from children with COVID-19 and healthy controls (**Data File S2**). We found significant enrichment of the NMF3: PD-L1+ ISG+ and the NMF6: ISG+ neutrophil subtypes in COVID-19 (**Fig. 2E**), plus a weak enrichment for the NMF5: G-MDSC subtype in COVID-19. Conversely, the NMF1: Pro-Neu signature was slightly enriched in the healthy controls as compared to pediatric COVID-19 (**Fig. 2E**). Taken together, these results confirm that acute COVID-19 infection in children induces an interferon-stimulated gene signature in neutrophils, similar to early stages of infection in adults (*1*).

### MIS-C is associated with a strong granulocytic myeloid-derived suppressor cell signature having similarities to severe COVID-19 in adults and sepsis

To define neutrophil activation in MIS-C, we performed differential expression analysis and gene set enrichment analysis comparing MIS-C samples with healthy pediatric controls (**Data File S2**). The most highly unregulated gene in MIS-C neutrophils was *GPR84*, a gene involved in ROS production and neutrophil activation (*18*) (**Fig. 3A**). Following closely was *ACER3*, a gene which has been shown to be upregulated in blood neutrophils of adult ARDS patients, several genes involved in glucose metabolism including *LDHA* and *ENO1*, and the complement C3 receptor *C3AR1* which may modulate neutrophil recruitment (*19*).

**Fig. 3.**
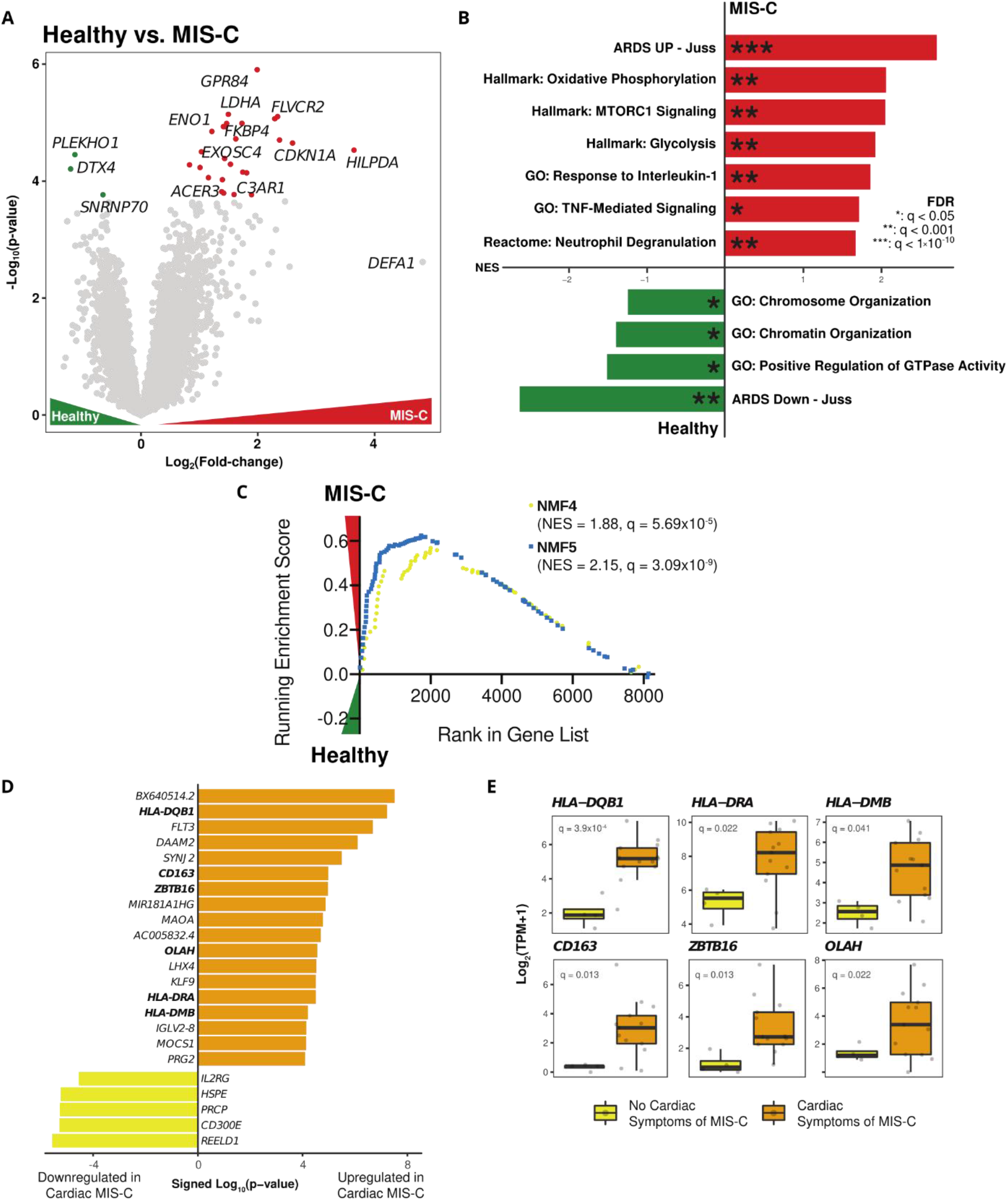
Neutrophils from MIS-C patients display a strong G-MDSC signature and are characterized by altered metabolism. **(A)** Volcano plot showing differentially expressed genes between MIS-C and pediatric healthy control samples. Color-coded points indicate genes that pass FDR correction with q < 0.05. **(B)** Gene set enrichment analysis for the differentially expressed genes in (A). Bar lengths correspond to normalized enrichment score (NES), with positive NES values indicating enrichment in MIS-C and negative values indicating enrichment in healthy controls. Asterisks denote significance as defined in the figure. **(C)** GSEA enrichment plots for the NMF4 (Immature) and NMF5 (G-MDSC) signatures, the two NMF neutrophil signatures which passed FDR correction. **(D)** Bar plots displaying the top differentially expressed genes between samples from MIS-C patients with and without cardiac involvement of disease. Bar length corresponds to the signed log10(p-value) for the differential expression analysis. Bolded genes are displayed individually in **(E)**. **(E)** Box plots of log2(TPM+1) values for selected genes differentially expressed in **(D)**. FDR q-values are from DESeq2.

The most highly enriched pathway in MIS-C was ARDS Up - Juss, a collection of genes which were found to be upregulated at least three-fold in adult ARDS blood neutrophils compared to healthy controls (*10*) (**Fig. 3B, Data File S2**). Accordingly, healthy control samples relative to MIS-C most prominently expressed the ARDS Down - Juss pathway, containing genes upregulated at least three-fold in healthy controls over ARDS samples. In agreement with the top differentially expressed genes, the next most highly enriched pathways in MIS-C were oxidative phosphorylation, glycolysis, and MTORC1 signaling, suggesting that MIS-C neutrophils are highly metabolically active. Finally, MIS-C samples were also enriched for neutrophil degranulation, TNF signaling, and IL-1 signaling, suggesting a mechanism by which neutrophils contribute to tissue damage in this highly inflammatory disease.

Next, we sought to determine which neutrophil subtypes were enriched in MIS-C. Gene set enrichment analysis revealed a strong enrichment of the NMF5: G-MDSC subtype in MIS-C, the subtype which was most strongly associated with severe disease and death in adult COVID-19 patients (*1, 2, 20*) (**Fig. 3C, Data File S2**), and NMF4, representing high levels of immature neutrophils that have been linked to increased NETosis, neutrophil degranulation, and activation (*1*).

We performed a direct comparison of MIS-C and acute pediatric COVID-19 neutrophils to validate our findings from the individual healthy control comparisons (**Data File S2**). Indeed, we observed a host of upregulated interferon-stimulated genes in acute COVID-19 that vanish in MIS-C, suggesting that the acute antiviral response has subsided in this late stage of disease, and instead we find several markers of increased cellular metabolism consistent with neutrophil activation (**Fig. S2A, Data File S2**). In agreement with these gene markers, the ISG+ neutrophil states (NMF3 and NMF6), were found to be highly enriched in acute pediatric COVID-19, while immature, NETosis-associated, and G-MDSC states (NMF1, NMF4, and NMF5, respectively), were enriched in MIS-C (**Fig. S2B)**. Additional markers for pediatric acute COVID-19 and MIS-C were highlighted in **Fig. S3**.

### MIS-C with cardiac involvement is associated with increased neutrophil antigen presentation and septic responses

Cardiac involvement is often associated with MIS-C but not required for diagnosis; several patients in our cohort had either myocarditis, ventricular dysfunction, or coronary arterial aneurysm, with two patients requiring ECMO. Thus, we next performed differential expression analysis to identify neutrophilic transcriptional differences between patients with (*n* = 13) or without (*n* = 4) cardiac symptoms of MIS-C (**Fig. 3D**). Several hallmark genes of sepsis and antigen presentation were enriched in cardiac MIS-C (**Fig. 3E**), including several of the MHC II genes (*HLA-DQB1*, *HLA-DRA*, *HLA-DMB)*. While neutrophils are not conventional antigen presenting cells, MHC II genes can be upregulated in PMNs by GM-CSF and IFNγ (*21*), and may compensate for decreased HLA-DR expression reported in conventional antigen-presenting cells (monocytes, DCs, and B cells) (*22*). In contrast to our findings in severe cardiac MIS-C, MHC II genes were found to be downregulated in neutrophils from adults with severe COVID-19 relative to adults with mild COVID-19. This could indicate a difference in the role of neutrophils between acute viral response in COVID-19 adults (*23*) and delayed onset antigenemia associated with MIS-C (*6*). Other markers of severe cardiac MIS-C are *CD163* which has been shown to be upregulated in PMNs of children with sepsis (*24*), *ZBTB16* which is high in adults with sepsis and severe COVID-19 (*1, 10*), and *OLAH* which is high in complex disease courses in sepsis (*25*).

### Neutrophils from children with MIS-C undergo high levels of spontaneous NETosis

As many of the highly upregulated genes and pathways from children with MIS-C are involved in neutrophil activation and potentially destructive neutrophil effector functions such as NET formation, granule release, and ROS production, we sought to visualize neutrophil activation by studying spontaneous NET formation by MIS-C neutrophils compared to healthy donor neutrophils. NETs were observed by microscopy using microfluidic technology; Neutrophils stimulated with 100 nM phorbol myristate acetate (PMA), a known, potent stimulator of NET formation, served as a positive control.

Neutrophils were isolated immediately after phlebotomy and plated without any added stimulants or neutrophil activators (experimental overview in **Fig. 4A**). While neutrophils from healthy pediatric controls showed minimal spontaneous NETosis over the four-hour period (**Movie S1**), neutrophils isolated from children with MIS-C showed high levels of spontaneous NETosis (percent of neutrophils undergoing NETosis: MIS-C 51.63±7.19% vs. healthy controls 0.17±0.21%, *P* < 0.0001; **Fig. 4, B and C**; **Movie S2**). The spontaneously formed NETs in MIS-C neutrophils were similar to PMA-stimulated NETs from healthy, pediatric children, reaching markedly elevated levels of NET formation (68.64±4.32% NETosis; **Fig. 4, B and C**; **Movies S2 and S3**).

**Fig. 4.**
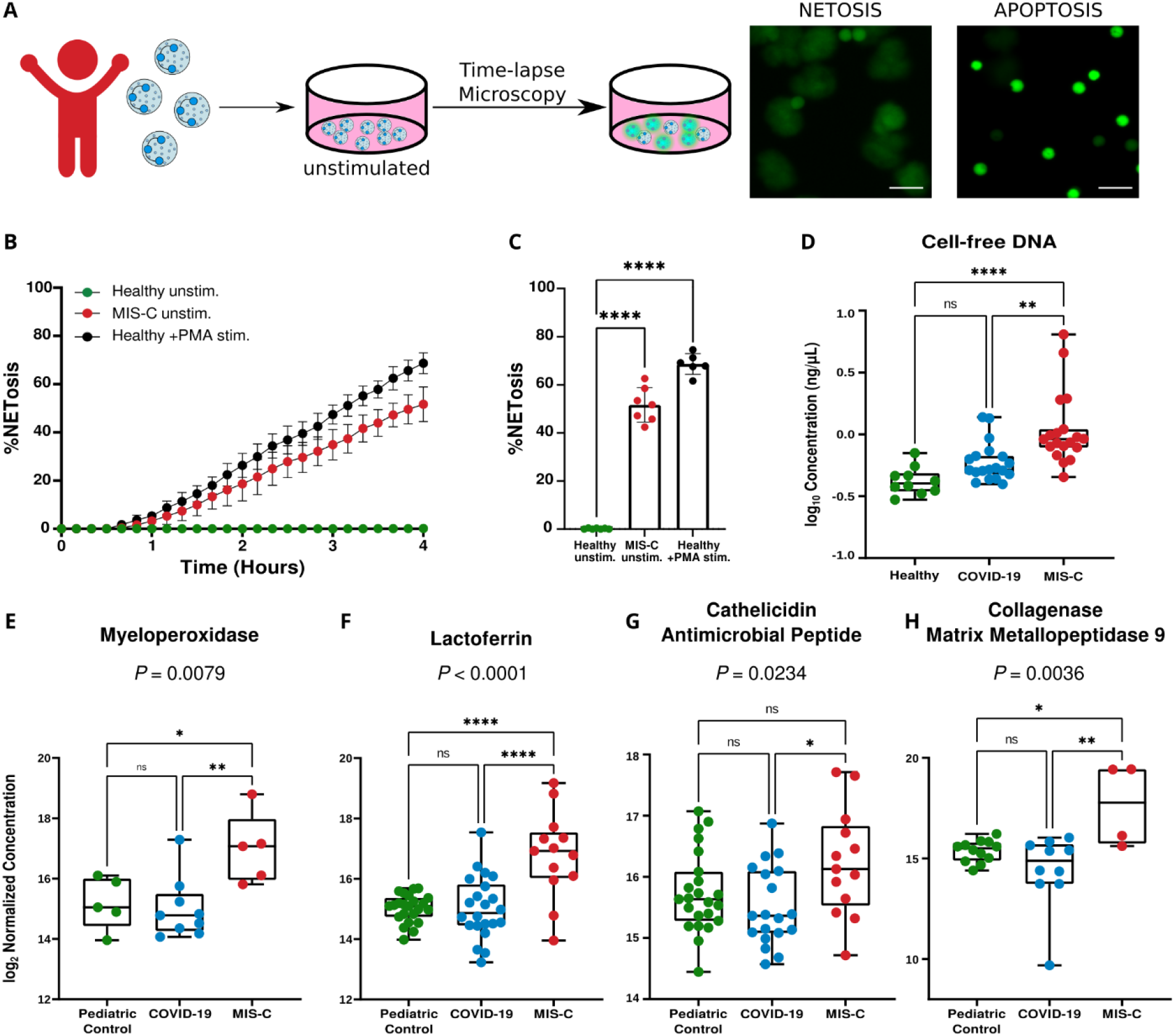
Fluorescence microscopy, cell-free DNA, and plasma protein markers implicate high levels of NETosis in MIS-C disease pathology. **(A)** Neutrophils were isolated from patients with MIS-C or healthy controls and plated on a 96-well plate in the absence of any stimulus to measure neutrophil activation via spontaneous NET release. Scale bar = 50 μM. **(B)** Temporal dynamics of NET release in unstimulated neutrophils from healthy children (*n* = 7) and children with MIS-C (*n* = 7), as well as healthy neutrophils stimulated with 100 nM PMA (*n* = 6). **(C)** End-point percentage of NETosis in unstimulated neutrophils from healthy children and children with MIS-C, and healthy neutrophils stimulated with 100 nM PMA. **(D)** Quantification of circulating cell-free DNA in plasma of healthy patients (*n* = 10), children with COVID-19 (*n* = 19), and children with MIS-C (*n* = 19). **(E), (F), (G), (H)** Peak values of myeloperoxidase **(E)**, lactoferrin **(F)**, cathelicidin antimicrobial peptide **(G)**, and collagenase matrix metallopeptidase 9 **(H)** quantified in plasma from pediatric controls (*n* = 23), children with acute COVID-19 (*n* = 19), and children with MIS-C (*n* = 13). Significance was determined by 1-way ANOVA with multiple comparisons. Mean values and standard deviation are presented. Statistical significance is defined as * *P* < 0.05, ** *P* < 0.01, *** *P* < 0.001, and **** *P* < 0.0001.

To corroborate evidence of NETosis in MIS-C, we probed for markers of NET release. We found children with MIS-C had significantly higher levels of cell-free DNA within their plasma (0.031±0.29 ng/μl) compared to children with acute COVID-19 (−0.22±0.16 ng/μl) (*P* = 0.002) and healthy children (−0.3757±0.11 ng/μl) (*P* < 0.0001) (**Fig. 4D**). Though cell-free DNA is nonspecific and could come from other forms of cell death, recent studies showed that circulating cell-free DNA in severe adult COVID-19 is primarily derived from hematopoietic cells, specifically neutrophils (*26*). We also used mass spectrometry to quantify levels of protein markers of NET release. In comparison to pediatric controls and children with COVID-19, children with MIS-C displayed high levels of neutrophil granule release, including significantly elevated levels of myeloperoxidase (MPO) (*P* = 0.0079), lactoferrin (LTF) (*P* < 0.0001), cathelicidin antimicrobial peptide (CAMP) (*P* = 0.0234), and matrix metallopeptidase 9 MMP9 (MMP9) (*P* = 0.0036) (**Fig. 4, E, F, G, and H**), supporting evidence of NET formation. Pediatric controls and children with acute COVID-19 had no detectable differences in these plasma markers of NET formation (**Fig. 4, E, F, G, and H**).

### Circulating anti-SARS-CoV-2 immune complexes stimulate NETosis

Current models of MIS-C pathogenesis suggest that the characteristic immune hyperactivation of the disease is driven by persistent antigenemia months after the initial SARS-CoV-2 infection (*6*). In one study, viral persistence in the GI tract led to the release of zonulin, a biomarker of intestinal permeability, which subsequently led to leakage of viral antigens into the bloodstream and subsequent immune hyperactivation (*6*). Thus, we hypothesized that SARS-CoV-2 antigen:antibody immune complexes (ICs) could be responsible for triggering NETosis in the vasculature, contributing to cardiovascular pathology seen in MIS-C. To probe this mechanism of increased NET formation in children with MIS-C, neutrophils isolated from healthy controls were treated with immune complexes that were generated by incubating patient-derived plasma with SARS-CoV-2 Spike antigen (experimental overview, **Fig. 5A, S4, and S5**). We observed low levels of NETosis when neutrophils were stimulated with Spike antigen plus buffer (12.54±3.08%; **Movie S4**) or plasma from non-COVID-19 healthy controls (13.70±3.94%; **Movie S5**), likely due to the absence of Spike-specific antibodies and ICs in these children (**Fig. 5, B and C**). We then stimulated neutrophils with convalescent COVID-19 plasma, in the absence of Spike antigen, to determine if SARS-CoV-2 antibodies alone could induce NETosis, and we again observed low levels of NETs in these stimulated neutrophils (13.31±1.86%; **Movie S6, Fig. 5, B and C**), indicating that neither antibodies nor antigen alone will induce NETosis.

**Fig. 5.**
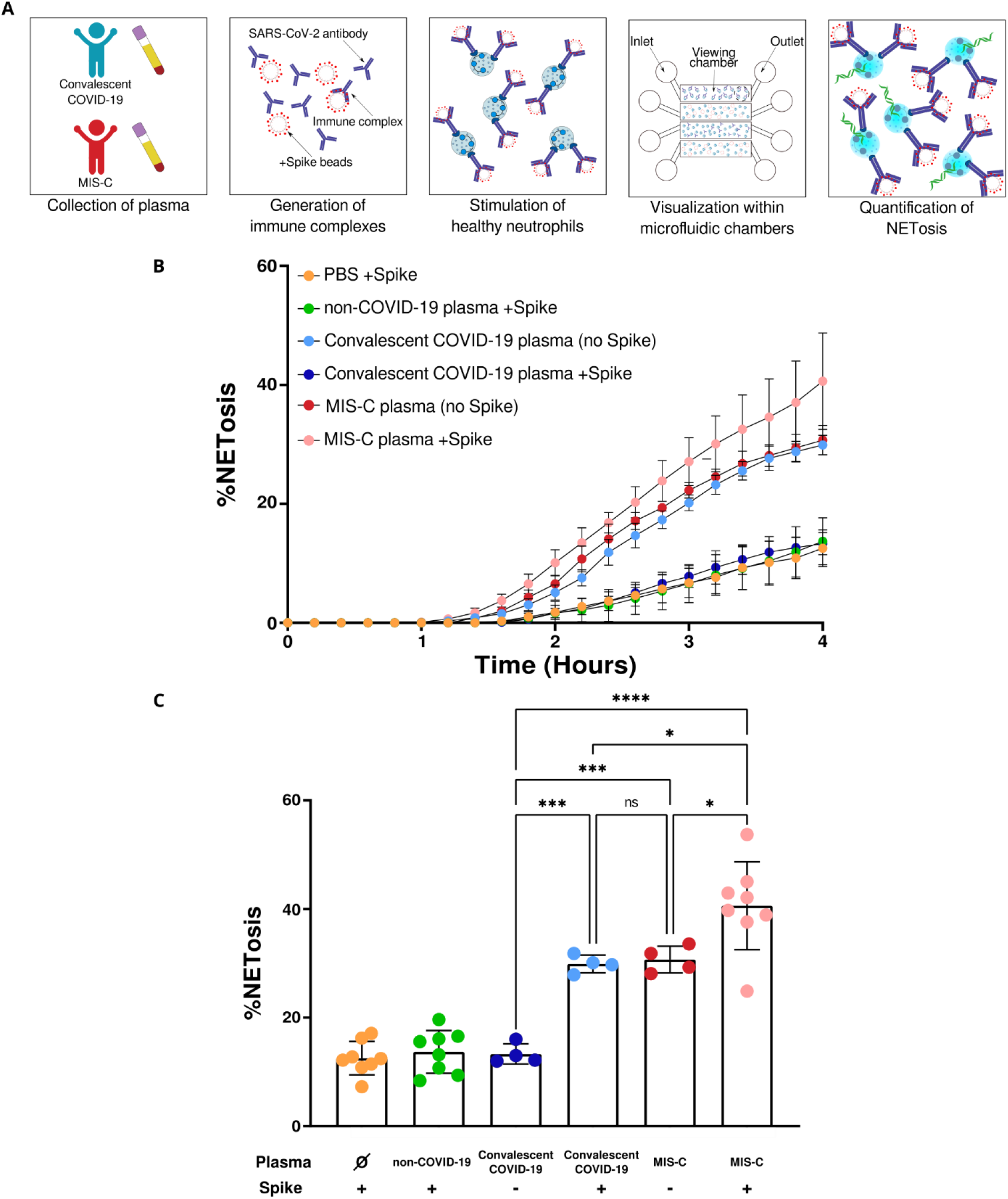
SARS-CoV-2 Spike immune complexes trigger NETosis. **(A)** Plasma was collected from children with MIS-C and convalescent COVID-19. Plasma was diluted 1:10 and incubated with beads coated with Spike proteins to generate immune complexes. Neutrophils were isolated from healthy children and stimulated with the immune complexes and visualized within 4 viewing channels of a microfluidic device. **(B)** Temporal dynamics of NET release in neutrophils stimulated with PBS-treated Spike beads (*n* = 8), non-COVID-19-plasma-coated Spike beads (*n* = 8), convalescent COVID-19 plasma (*n* = 4), convalescent COVID-19-plasma-coated Spike beads (*n* = 4), MIS-C plasma (*n* = 4), MIS-C-plasma-coated Spike beads (*n* = 8). **(C)** End-point percentage of NET release in neutrophils stimulated with PBS-treated Spike beads (*n* = 8), non-COVID-19-plasma-coated Spike beads (*n* = 8), convalescent COVID-19 plasma (*n* = 4), convalescent COVID-19-plasma-coated Spike beads (*n* = 4), MIS-C plasma (*n* = 4), MIS-C-plasma-coated Spike beads (*n* = 8). Significance was determined by 1-way ANOVA with multiple comparisons. Mean values and standard deviation are presented. Statistical significance is defined as * *P* < 0.05, ** *P* < 0.01, *** *P* < 0.001, and **** *P* < 0.0001.

However, the addition of Spike antigen to convalescent COVID-19 plasma, allowing formation of SARS-CoV-2 specific antibody:antigen ICs, induced significant NETosis (29.89±1.62%) (**Movie S7**) in comparison to convalescent COVID-19 plasma alone (*P* = 0.0005) (**Fig. 5, B and C**). This level of NETosis was comparable to that seen when healthy neutrophils were stimulated with MIS-C plasma (30.69±1.46%; **Movie S8**). Adding additional Spike antigen to MIS-C plasma stimulated an even greater amount of NET formation (40.61±8.10%; **Fig. 5, B and C**; **Movie S9**). These results indicate that SARS-CoV-2 antigenemia in the setting of a mature humoral response appears to instigate excessive neutrophil activation with hyperinflammatory and detrimental intravascular NET formation in MIS-C.

## DISCUSSION

In this study, we report the first in-depth characterization of neutrophils in pediatric acute COVID-19 and MIS-C. Utilizing neutrophil bulk RNA-seq, plasma protein expression, and functional analyses of neutrophils from children with acute COVID-19, MIS-C, and pediatric controls, we highlight major differences in the neutrophil response that help to explain the divergent presentations of these SARS-CoV-2-related diseases.

MIS-C, though SARS-CoV-2-related, presents as a completely distinct set of symptoms and neutrophil phenotypes from acute COVID-19 in both children, shown here, and previous reports in adults (*1*). This is likely driven by the absence of viremia in MIS-C (*27*), and instead may be linked to SARS-CoV-2 antigenemia without replication-competent virus in the setting of a mature humoral response. Studies have demonstrated that viral RNA is still detected in the stool of children weeks after infection or exposure (*6*), perhaps through a mechanism of antibody-dependent enhancement (ADE) in which non-neutralizing antibodies facilitate uptake of virus into ACE2-negative Fc-receptor-positive cells and allow for continued replication (*28*). Persistence of virus is associated with increased gut permeability through the release of zonulin by gut epithelial cells (*6*), allowing viral antigens to escape into the bloodstream, where primed antibodies await. The formation of immune complexes triggers a hyperinflammatory immune response with neutrophil activation and NET formation within the vasculature (**Fig. 6**).

**Fig. 6.**
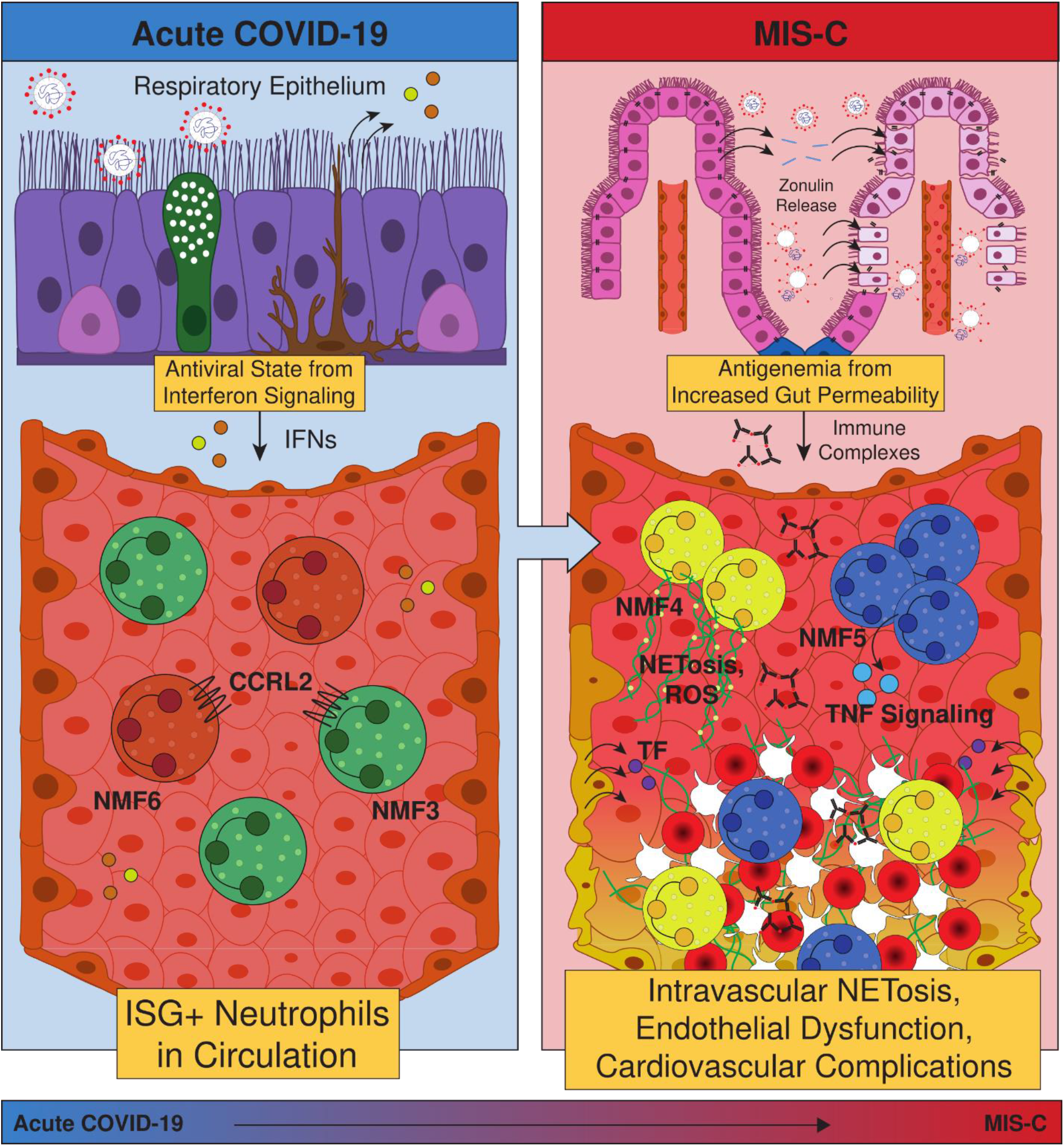
Schematic of the role of neutrophils in pediatric COVID-19 and MIS-C. Acute SARS-CoV-2 infection begins in the respiratory epithelium, and interferon signaling induces ISG+ neutrophils in circulation. Weeks later, viral persistence in the gut lumen results in zonulin release from gut epithelial cells, leading to loss of tight junctions and SARS-CoV-2 antigen leakage. Finally, immune complexes trigger NETosis and induce G-MDSCs, resulting in endothelial dysfunction and cardiovascular complications.

As a result of this distinct mechanism, neutrophils in MIS-C are characterized by upregulated genes and pathways associated with G-MDSCs and the formation of NETs, which is highly distinct from the antiviral gene signature seen in acute pediatric COVID-19. Neutrophils in patients with MIS-C more closely resemble those found in patients with non-viral ARDS (*10*) or sepsis (*20*) rather than mild acute COVID-19 in that they appear to take on an immature, T cell-suppressive phenotype. This activated and destructive phenotype is hypothesized to cause the systemic damage seen in patients with MIS-C, with potential mechanisms being the interaction of neutrophil Fc receptors with SARS-CoV-2 antigen immune complexes, or TNFɑ signaling via NF-κB which in excess can dysregulate T cells (*29*) or activate further NETosis. Indeed, increased NF-κB signaling has been shown to be upregulated in monocytes and dendritic cells in MIS-C, especially in severe cardiac MIS-C (*30*), but prior reports have not examined neutrophils. Activated neutrophils have been implicated in the pathology of myocarditis through infiltration into the heart, with evidence of NETs have been found in endomyocardial biopsies from patients with myocarditis (*31–33*). As activated neutrophils and viral particles have been found in the myocardium post-mortem in MIS-C patients (*34, 35*), increased antigen presentation by neutrophils and upregulation of septic responses by neutrophils may play a role in pathology driving severe, cardiac complications of MIS-C.

In this study, we also provide the first visual evidence of high rates of spontaneous NETosis in neutrophils from children with MIS-C, with imaging experiments performed within hours of fresh whole blood collection. Furthermore, we provide mechanistic insight into the abundant NET formation by showing that SARS-CoV-2 antigen:patient-derived antibody immune complexes are capable of triggering NETosis in healthy donor neutrophils. The observed NETosis has several implications for the disease pathology observed in MIS-C, because although NETs are effective at capturing and destroying pathogens at mucosal barriers, they can be a source of excessive inflammation and coagulation when released in circulation. First, NET components such as MPO can generate ROS and can directly damage endothelium through several mechanisms, leading to the release of tissue factor (TF). NETs themselves often contain TF (*36*); thus, both NETs and the damaged endothelium can initiate the extrinsic coagulation cascade. Second, the abundant extracellular DNA can serve as a damage-associated molecular pattern (DAMP) (*37*), triggering additional inflammation alongside inflammatory cytokine production, and the negative charge can initiate the intrinsic coagulation cascade (*38*). Taken together, these mechanisms can serve as a scaffold for platelets, induce hypercoagulability and endothelial dysfunction, and lead to thrombosis, vasculitis, and cardiovascular complications, providing a uniting framework for MIS-C pathogenesis (**Fig. 6**).

Neutrophil activation in MIS-C starkly contrasts the interferon-stimulated gene profile seen in neutrophils from children with acute COVID-19. In children, the antiviral response to SARS-CoV-2 is sufficient to clear acute infection without severe symptoms, as evidenced by the mild or asymptomatic presentations in most cases. While live, infectious SARS-CoV-2 can be detected in the upper airways of children (*39*) accounting for the antiviral response, there is little to no viremia detected in pediatric COVID-19 (*27*). This picture may be similar to that seen in adults with mild COVID-19. However, in adults with moderate to severe disease, impaired antiviral signaling can lead to viremia in a larger fraction of cases, resulting in a more severe presentation of acute COVID-19 in these individuals (*40–42*). Thus, containment of the virus in the upper respiratory tract in children likely prevents the progression to more severe circulating neutrophil phenotypes seen in life-threatening COVID-19 in adults, and it is unlikely that circulating neutrophils alone determine disease severity in acute pediatric COVID-19.

There is an increasing appreciation of neutrophil activation in severe COVID-19 in adults, and therapies are being designed to target neutrophil hyperactivation. However, in most children, their immune system is already able to manage SARS-CoV-2 due to a strong interferon-mediated antiviral response (*12, 13*). In both adults and children, SARS-CoV-2 viremia or antigenemia and the resultant immune complexes appear to be responsible for deleterious neutrophil extracellular trap formation. While therapies blunting excessive neutrophil activation could improve clinical outcomes in severe disease, mitigating the antigenemia or viremia will be critical. Previous studies have indicated that preventing SARS-CoV-2 antigen leakage from the gut with the zonulin antagonist, larazotide, may be effective in preventing antigenemia-included hyperinflammation in MIS-C (*6*). While vaccination and avoidance of infection are the best prevention strategies, if SARS-CoV-2 infection does occur and MIS-C ensures, future studies may explore combination therapies that both prevent antigenemia by enhancing tight junctions in gut epithelium and simultaneously inhibit systemic NET formation in order to improve upon the current treatment paradigm.

We acknowledge several limitations of our study. First, we utilized bulk transcriptomics due to the technical challenges of performing single-cell RNA-sequencing on neutrophils, so each sample represents a mixture of neutrophils with distinct gene expression programs. However, our high purity double isolation method left little contamination with other cell types. Second, we acknowledge the small size of the cohort, although it represents the largest cohort of its kind due to the low incidence rate of the disease. Third, based on the sample size limitation we were unable to define new gene expression states that may have been specific to pediatric neutrophils and had to rely on the adult-derived signatures, though that work demonstrated a relatively small set of conserved neutrophil states across many different disease contexts. Fourth, MIS-C can present heterogeneously and there was a mixture of treatment regimens and duration of disease among the patients, which may have resulted in greater heterogeneity in the gene expression profiles. All limitations considered, numerous technical challenges and time-sensitive issues were overcome in this study to allow for deep characterization of neutrophil profiles in acute pediatric COVID-19 and MIS-C.

In summary, our study provides much needed mechanistic insight into the drivers of pathology in MIS-C and its departure from mild acute SARS-CoV-2 infection in children. We link existing knowledge of gut mucosal breakdown and antigenemia in MIS-C to increased immune complex-dependent NETosis and a shift away from the antiviral state towards a sepsis-like G-MDSC phenotype, highlighting potential connections between neutrophil activation and the characteristic cardiovascular pathology to suggest pathways of intervention to improve the standard of care.

## MATERIALS AND METHODS

### Study Design

Biospecimens were obtained from pediatric patients at Massachusetts General Hospital (MGH) under the institutional review board (IRB) approved ‘MGH Pediatric COVID-19 Biorepository’ (no. 2020P000955). Healthy pediatric controls were collected under the IRB-approved ‘the Center for Celiac Research Pediatric Biorepository’ (no. 2016P000949). Informed consent, and assent when appropriate, were obtained in accordance with IRB guidelines from patients and/or parents/guardians before study enrollment. Patients classified as COVID-19 positive had positive SARS-CoV-2 RT-PCR results upon enrollment. Patients classified as MIS-C were diagnosed per CDC guidelines.

### Neutrophil isolation and lysis

Blood samples were processed as previously described (*43*). Briefly, blood samples were collected in vacutainer tubes containing Ethylenediaminetetraacetic Acid (EDTA) anticoagulant. After plasma collection, PBMCs were isolated and collected via Ficoll density gradient. Neutrophils were isolated from the remaining blood pellet via negative selection using the EasySep Direct Neutrophil Isolation Kit (STEMCELL Technologies, Cat# 19666) following manufacturer’s directions. 1×10^5^ cells were centrifuged at 300g for 5 minutes at room temperature with brakes activated. Neutrophils were lysed with 100 μl of RNA lysis buffer (TCL) (Qiagen, Cat# 1031576) with 1% β-mercaptoethanol and frozen at −80°C.

### Validation of isolated neutrophil purity

Nucleated cells were collected from whole blood via red blood cell sedimentation using HetaSep solution (STEMCELL Technologies, Cat# 07906). Neutrophils were isolated as previously described (plasma centrifuged, PBMCs isolated and collected, and neutrophils isolated from remaining red blood cell pellet). Isolated neutrophils and nucleated cells were centrifuged at 300g for 5 minutes at room temperature with the brakes off. The cells were then resuspended in 50 μl of FACS buffer (0.5% BSA in PBS) and then fixed in 1% paraformaldehyde in PBS. All fixed cells were then stained with CD66b (1:200 dilution) (Biolegend, Cat# 305103), CD14 (1:20 dilution) (Biolegend, Cat# 325615), and DRAQ5 (1:2000 dilution) (Cell Signaling Technology, Cat# 4084S). Data was obtained through the Amnis ImageStreamX Mark II imaging flow cytometer and INSPIRE software (EMD Millipore, Billerica, MA, USA). The accompanying IDEAS software (EMD Millipore) was used to perform data analysis.

### Patient matched plasma isolation

Blood samples were processed as previously described (*43*). Briefly, blood samples were collected in EDTA vacutainer tubes. Plasma was collected by centrifuging tubes at 1000g for 10 minutes at room temperature with brakes activated. Aliquoted plasma was stored at −80°C.

### Smart-Seq2 cDNA preparation

cDNA was prepared from bulk populations of 1×10^5^ neutrophils as recently described (*1*) using a modified reverse transcription step (*44*). Neutrophil lysates were thawed on ice, and 20 μl per sample was added to a 96-well plate prior to brief centrifugation. Following RNA purification with Agencourt RNAClean XP SPRI beads (Beckman Coulter, Cat# A63987), samples were resuspended in 4 μl of Mix-1 [Per 1 sample: 1 μl (10 μM) RT primer (DNA oligo) 5′– AAGCAGTGGTATCAACGCAGAGTACT30VN-3′; 1 μl (10 μM) dNTPs; 1μl (10%, 4 U/μl) recombinant RNase inhibitor; 1 μl nuclease-free water], denatured at 72°C for 3 minutes and immediately cooled on ice for 1 minutes. Next, 7 μl of Mix-2 [Per 1 sample: 0.75 μl nuclease-free water; 2 μl 5X RT buffer (Thermo Fisher Scientific, Cat# EP0753); 2 μl (5 M) betaine; 0.9 μl (100 mM) MgCl2; 1 μl (10 μM) TSO primer (RNA oligo with LNA) 5′-AAGCAGTGGTATCAACGCAGAGTACATrGrG+G-3′; 0.25 μl (40 U/μl) recombinant RNase inhibitor; 0.1 μl (200 U/μl) Maxima H Minus Reverse Transcriptase] was added. Reverse transcription was performed at 50°C for 90 minutes, followed by a 5 minute incubation at 85°C. Next, 14 μl of Mix-3 [Per 1 sample: 1 μl nuclease-free water; 0.5 μl (10 μM) ISPCR primer (DNA oligo) 5′-AAGCAGTGGTATCAACGCAGAGT-3′; 12.5 μl 2X KAPA HiFi HotStart ReadyMix] was added to each well and amplification was performed at 98°C for 3 minutes, followed by 16 cycles of [98°C for 15 s, 67°C for 20 s, and 72°C for 6 minutes], and final extension at 72°C for 5 minutes. Primer removal and cDNA purification was performed using AgencourtAMPureXP SPRI beads (Beckman Coulter, Cat# A63881). Concentration measurements were determined with the Qubit dsDNA high sensitivity assay kit (Invitrogen, Cat# Q32854) on the Cytation 5 Microplate Reader (BioTek), and cDNA size distribution was measured using the High-Sensitivity DNA Bioanalyzer Kit (Agilent, Cat# 5067-4626).

### Library construction and sequencing

Libraries were constructed with the Nextera XT Library Prep kit (Illumina, Cat# FC-131-1024) using custom indexing adapters (*44*) in a 384-well PCR plate. Residual primer dimers were removed with an additional cleanup step. One pooled library containing 26 samples were sequenced on a NextSeq 500 sequencer (Illumina) using paired-end 38-base reads. Following quality control and acquisition of additional samples, a second pooled library containing 40 samples was sequenced using the same method.

### Spontaneous NET release fluorescence microscopy data acquisition

Blood was processed from healthy pediatric patients and MIS-C patients as previously described (*43*). From the isolated neutrophils, an aliquot of 1×10^4^ cells in 50 μl RPMI (no FBS) were collected. Neutrophils were plated in a single well of a black, clear bottom 96-well plate. For the unstimulated neutrophils, 50 μl of RPMI was added to the well. For the stimulated neutrophils, 50 μl of phorbol myristate acetate (PMA) (Sigma Aldrich, Cat# P1585) was added to a final concentration of 100 nM. 50 μl of SYTOX green (Thermo Fisher, Cat# S7020) in RPMI was added to a final concentration of 2 μM. NETosis was imaged using the Personal AUtomated Lab Assistant (PAULA) Cell Imager (Leica Microsystems) placed in a 37°C incubator with 5% CO_2_. Brightfield and FITC images were taken every 10 minutes for 4 hours to allow for temporal imaging of NET formation.

### Cell-free DNA quantification

cfDNA quantification was performed using the Qubit dsDNA High Sensitivity Assay Kit (Invitrogen, Cat# Q32854) according to the manufacturer’s protocol. 98 μl of Qubit DNA dye was aliquoted per well of a 96-well black clear bottom plate (Corning, Cat# 3904). 2 μl of plasma sample was added to each well of the assay plate, and fluorescence was measured on a Cytation 5 Microplate reader at 523 nm.

### Plasma protein measurement

Samples were prepared as previously described(*6*). 100 μg were denatured, reduced, alkylated and purified using solid-phase-enhanced sample-preparation (SP3) technology prior to lysc and tryptic digest (*45, 46*). 25 μg of the resulting peptides were subsequently labeled using TMTpro reagents (Thermo Scientific) according to manufacturer’s instructions (*47, 48*). Labeled samples were combined and fractionated using a basic reversed phase HPLC (*45*). The fractions were analyzed via reversed phase LC-M2/MS3 on either an Orbitrap Fusion Lumos or an Orbitrap Eclipse mass spectrometer using the Simultaneous Precursor Selection (SPS) supported MS3 method (*49, 50*) essentially as described previously (*51*) and using Real-Time Search when using the Orbitrap Eclipse (*52*). MS2 spectra were assigned using a SEQUEST-based (*53*) in-house built proteomics analysis platform (*54*) using a target-decoy database-based search strategy to assist filtering for a false-discovery rate (FDR) of protein identifications of less than 10% (*55*). Searches were done using the Uniprot human protein sequence database (UP000005640) (*55*) using an in-house-built platform (*53*). Search strategy included a target-decoy database-based search in order to filter against a false-discovery rate (FDR) of protein identifications of less than 1% (*55*). An average signal-to-noise value of larger than 10 per reporter ion as well as with an isolation specificity (*50*) of larger than 0.75 were considered for quantification. Lysozyme, Myeloperoxidase, Gelatinase matrix metallopeptidase 9, Lactoferrin, and Cathelicidin antimicrobial peptide were quantified. Protein concentration data were normalized as previously described (*47*). A 1-way ANOVA was performed to compare relative protein concentrations between COVID-19, MIS-C, and pediatric control samples. If the ANOVA showed significance with a *P* value of less than 0.05, a 2-sided *t* test was performed on paired sample groups to determine which proteins varied significantly between cohorts.

### Design and fabrication of the microfluidic devices

Microfluidic devices used for time lapse imaging of NETosis were designed using AutoCAD (Autodesk) and fabricated using standard photolithographic and soft lithography approaches **(Fig. S4, A and B)**. Briefly, a master negative mold was prepared by spin-coating a silicon wafer with negative photoresist (SU-8, Microchem, MA) to a thickness of 100 μm. The design was patterned onto the wafer by exposure to UV light through a mylar mask (Fineline Imaging, CO) and developing away the unexposed areas. The silicon wafer was then used as a template for casting PDMS devices. PDMS base and primer were mixed at a ratio of 10:1 and poured over the wafer, degassed for at least 1 hour, then baked overnight at 65°C. Inlets and outlets were punched using a 1 mm diameter biopsy punch (Harris UniCore, Millipore Sigma), and the entire device liberated from the PDMS block using an 8 mm diameter punch. PDMS devices were treated with oxygen plasma and bonded irreversibly to oxygen plasma treated glass-bottom well plates (Mattek, MA) by heating to 75°C for 10 minutes.

### Immune complex generation

Immune complexes were generated as previously described (*56*). Briefly, SARS-CoV-2 Spike protein were biotinylated and conjugated to NeutrAvidin beads and incubated with 1:10 diluted plasma samples.

### NETosis assay

10 μl of RPMI was loaded into the inlet port of the microfluidic device until a droplet was formed on the outlet port. RPMI was added to the wells until the devices were submerged. The plate was placed in a 37°C, 5% CO_2_ incubator until the neutrophil solution was prepared to be loaded. Neutrophils were isolated from a healthy patient cohort. 1-2 ml whole blood was collected in EDTA vacutainer tubes and neutrophils were isolated via negative selection using the Easysep Direct Neutrophil Isolation Kit following manufacturer’s directions. Isolated neutrophils were stained with 32 μM Hoechst 3342 dye for 10 minutes at 37°C, 5% CO_2_. Stained neutrophils were then resuspended in RPMI (no FBS) to a concentration of 1×10^7^ cells/ml. 10 μl of cells were mixed with 10 μl of stimulant, and 10 μl of SYTOX green in RPMI (no FBS) to a final concentration of 2 μM. 5 μl of this solution was loaded into the inlet port of the microfluidic device using a standard pipette tip until cells were observed to exit the outlet port. Brightfield and FITC fields were imaged at 10x magnification using a fully automated fluorescent Nikon TiE inverted wide field microscope with a biochamber heated to 37°C with 5% CO_2_. Each device was imaged every 12 minutes for 4 hours to allow for temporal imaging of NET formation. An initial image was taken with brightfield and DAPI at the beginning of the experiment to count fluorescent stained neutrophils.

## QUANTIFICATION AND STATISTICAL ANALYSIS

### RNA-seq alignment

Raw FASTQ files were aligned to a custom genome using the Terra platform of the Broad Institute. Alignment was performed using STAR v2.5.3a, and expression quantification was performed using RSEM v1.3.0. The custom FASTA was generated from the Homo sapiens genome assembly GRCh38 (hg38, excluding ALT, HLA, and Decoy contigs according to the Broad Institute GTEx-TOPMed RNA-seq pipeline, https://github.com/broadinstitute/gtex-pipeline/) with an appended SARS-CoV-2 genome. GENCODE v35 with the appended SARS-CoV2 GTF was used for annotation.

### Quality control

RNA-SeQC 2 (*57*) (https://github.com/getzlab/rnaseqc) was used to calculate quality control metrics for each sample. Samples were excluded if they did not meet the following criteria: 1) percentage of mitochondrial reads less than 20%, 2) greater than 10,000 genes detected with at least 5 unambiguous reads, 3) median exon CV less than 1, 4) exon CV MAD less than 0.75, 5) exonic rate greater than 25%, 6) median 3’ bias less than 90%. One sample was excluded for having low neutrophil content and high B cell contamination as estimated by CIBERSORTx, described in the next section. Only one sample was kept per patient: if more than one passed quality control, the earlier or pre-treatment sample was selected. Thus, out of 55 sequencing reactions, 48 samples were carried forward for analysis. Genes were included in the analysis if they were expressed at a level of 0.1 TPM in at least 20% of samples and if there were at least 6 counts in 20% of samples. In total, 15406 genes passed the filtration criteria.

### Neutrophil fraction estimation and contamination control

CIBERSORTx (*15*) was used to estimate total neutrophil, mature neutrophil, immature neutrophil, T/NK cells (either T or NK, due to similar gene expression profiles), B, plasmablast, and monocyte content in each sample. We generated a signature matrix from the single-cell RNA-seq data of PBMCs and neutrophils from whole blood from Cohort 2 of the Schulte-Schrepping et al dataset (*2*) using the pseudobulking method previously described (*1*). CIBERSORTx was run using default parameters.

### Dimensionality reduction and visualization

PCA and UMAP were performed in R using prcomp() and umap() with default parameters.

### Differential expression analysis

Differential expression analyses were performed using the DESeq2 package in R (*58*) without additional covariates.

### Gene set enrichment analysis

Gene set enrichment analysis was performed using the fgsea package in R using MSigDB Release v7.2 pathways from the H, C5 GO BP databases. We also added custom MSigDB pathways using the keyword “neutrophil”, and the gmt file is available in the Github. In addition, we added gene sets defining neutrophil states in ARDS (*10*), blood and lung tissue of lung cancer patients (*17*), sepsis (*20*), single-cell neutrophil clusters in COVID-19 (*2*), and bulk RNA-seq signatures of neutrophils in COVID-19 (*1*).

### Sample Pathway Scoring

Bulk RNA-seq samples were scored according to their expression of NMF neutrophil state gene signatures according to a previously described method used to control for sample complexity (*59*). We defined the pathway score for each sample as the average expression of the genes in the gene set minus the average expression of genes in a control gene set. Genes were ranked according to average expression across all samples and divided into 25 bins, and for each gene in the NMF signature, 100 different genes were selected from the same bin to create a control gene set with comparable expression levels which is 100-fold larger.

### Disease marker gene selection and heatmap

To identify genes uniquely associated with acute pediatric COVID-19 or MIS-C, we performed three differential expression analyses: MIS-C vs. COVID-19 and controls, COVID-19 vs. MIS-C and controls, and controls vs. MIS-C and COVID-19. We filtered the results for p_adj_ > 0.05 and only selected positive markers (log(fold-change) greater than 0) and capped the lists at 50 genes. There were no significant marker genes for control neutrophils. Heatmaps of the gene markers for the two diseases were generated with the pheatmap package in R. Row (genes) are ordered according to P value, and columns are ordered according to the clinical subtypes shown in the legend.

### Quantification of NETosis

Detection of NETs was performed automatically on the FITC channel using the TrackMate plugin in FIJI (*60, 61*). As neutrophils undergo NETosis, the nuclear membrane of the cell breaks down, allowing for SYTOX green to stain the DNA and to be visualized via the FITC channel. NETs are quickly dispersed, and the fluorescence is diffused. Apoptotic cells, on the other hand, can become stained with SYTOX green as the membrane breaks down (**Fig. S5A**), however the fluorescence signal is stable and long lasting (**Fig. S5B**). Methodology was validated using manual tracking within FIJI (*60*) to determine average length of time of cells undergoing NETosis against cells undergoing apoptosis, where a cutoff differentiating the two was defined at 1.5 hours (**Fig. S5C**). Thus, NETs were defined as a tracked cell with a duration ≤ 1.5 hours and apoptotic cells were defined as tracked cells with a duration > 1.5 hours. Total number of neutrophils was counted using the DAPI channel at time *t* = 0 (*60*). Percentage of NETs was calculated using the number of NETs released at a specific time point divided by the total number of neutrophils.

### Statistical analysis

To compare multiple groups, results were compared by 1-way ANOVA with multiple comparisons in GraphPad Prism v9. Mean values and standard deviation are presented. Statistical significance is defined as * *P* < 0.05, ** *P* < 0.01, *** *P* < 0.001, and **** *P* < 0.0001, unless otherwise noted in the figure.

## Supplementary Materials

**Fig. S1**. Neutrophil isolation and quality control for purity, related to Fig. 1.

**Fig. S2**. Direct comparison of acute pediatric COVID-19 and MIS-C, related to Fig. 3.

**Fig. S3**. Heat map comparison of healthy children, acute pediatric COVID-19, and MIS-C, related to Fig. 3.

**Fig. S4**. Design and schematic of microfluidic device, related to Fig. 5.

**Fig. S5**. Methodology and validation of quantification of NETosis, related to Figs. 4 and 5.

**Table S1**. Demographics and clinical characteristics of pediatric acute COVID-19 patients included in the study.

**Table S2**. Demographics and clinical characteristics of pediatric MIS-C patients included in the study.

**Data File S1**. Neutrophil RNA-seq quality control information.

**Data File S2**. Differential expression and GSEA results.

**Data File S3**. Gene expression matrices in Counts and TPM.

**Movie S1**. Spontaneous NETosis of neutrophils isolated from a healthy pediatric patient captured by fluorescence microscopy.

**Movie S2**. Spontaneous NETosis of neutrophils isolated from a child with MIS-C captured by fluorescence microscopy.

**Movie S3**. NETosis of neutrophils isolated from a healthy pediatric patient stimulated with 100 nM PMA captured by fluorescence microscopy.

**Movie S4**. NETosis of isolated neutrophils isolated from a healthy patient stimulated with PBS-treated beads captured by fluorescence microscopy.

**Movie S5**. NETosis of isolated neutrophils isolated from a healthy patient stimulated with non-COVID-19 plasma-treated Spike beads captured by fluorescence microscopy.

**Movie S6**. NETosis of isolated neutrophils isolated from a healthy patient stimulated with convalescent COVID-19 plasma captured by fluorescence microscopy.

**Movie S7**. NETosis of isolated neutrophils isolated from a healthy patient stimulated with convalescent COVID-19 plasma-treated Spike beads captured by fluorescence microscopy.

**Movie S8**. NETosis of isolated neutrophils isolated from a healthy patient stimulated with MIS-C plasma-treated captured by fluorescence microscopy.

**Movie S9**. NETosis of isolated neutrophils isolated from a healthy patient stimulated with MIS-C plasma-treated Spike beads captured by fluorescence microscopy.

## Acknowledgments

We would like to acknowledge the efforts of all members of the MGH Pediatric COVID-19 Biorepository, especially Ms. Madeleine Burns, Ms. Eva Farkas, and Ms. Christina Lee, and the Center for Celiac Research Pediatric Biorepository, especially Dr. Maureen M. Leonard and Ms. Victoria Kenyon for their assistance in recruiting and processing biospecimens for the study.

## Funding

National Heart, Lung, and Blood Institute grant 5K08HL143183 (LMY)

Massachusetts General Hospital Executive Committee on Research, COVID-19 Clinical Trials Initiative grant (LMY)

Department of Pediatrics at Massachusetts General Hospital *for* Children (LMY)

## Author contributions

Conceptualization: MSF, LMY

Methodology: BPB, TJL, YB, FE, MEL, JPD, ALKG, SH, SP, WH, GA, DI, MSF, LMY

Validation: BPB, TJL, YB, FE, MEL, JPD, ALKG, SH, SP, WH, GA, DI, MSF, LMY

Formal Analysis: BPB, TJL, YB, FE, SH, GA, DI, MSF, LMY

Investigation: BPB, TJL, YB, FE, MEL, JPD, ALKG, SH, SP, WH, AE, AF, GA, DI, MSF, LMY

Resources: SP, WH, AE, AF, GA, DI, MSF, LMY

Data Curation: BPB, TJL, SH, MSF, LMY

Writing- Original Draft: BPB, TJL, MSF, LMY

Writing- Review and Editing: BPB, TJL, YB, FE, MEL, JPD, ALKG, SH, SP, WH, AE, AF, GA, DI, MSF, LMY

Visualization: BPB, TJL, FE, MEL, DI, MSF, LMY

Supervision: MSF, LMY

## Data and materials availability

All data are available in the main text or the supplementary materials. The read count matrix and TPM matrix used in this study will be available in GEO and **Data File S3**. All code used for the analysis is deposited in GitHub at https://github.com/lasalletj/Pediatric_COVID_MISC_Neutrophils.

## References and Notes

1. T. J. LaSalle, A. L. K. Gonye, S. S. Freeman, P. Kaplonek, I. Gushterova, K. R. Kays, K. Manakongtreecheep, J. Tantivit, M. Rojas-Lopez, B. C. Russo, N. Sharma, M. F. Thomas, K. M. Lavin-Parsons, B. M. Lilly, B. N. McKaig, N. C. Charland, H. K. Khanna, C. L. Lodenstein, J. D. Margolin, E. M. Blaum, P. B. Lirofonis, A. Sonny, R. P. Bhattacharyya, B. A. Parry, M. B. Goldberg, G. Alter, M. R. Filbin, A. C. Villani, N. Hacohen, M. Sade-Feldman, Longitudinal characterization of circulating neutrophils uncovers distinct phenotypes associated with disease severity in hospitalized COVID-19 patients. bioRxiv, (2021).

2. J. Schulte-Schrepping, N. Reusch, D. Paclik, K. Baßler, S. Schlickeiser, B. Zhang, B. Krämer, T. Krammer, S. Brumhard, L. Bonaguro, E. De Domenico, D. Wendisch, M. Grasshoff, T. S. Kapellos, M. Beckstette, T. Pecht, A. Saglam, O. Dietrich, H. E. Mei, A. R. Schulz, C. Conrad, D. Kunkel, E. Vafadarnejad, C. J. Xu, A. Horne, M. Herbert, A. Drews, C. Thibeault, M. Pfeiffer, S. Hippenstiel, A. Hocke, H. Müller-Redetzky, K. M. Heim, F. Machleidt, A. Uhrig, L. Bosquillon de Jarcy, L. Jürgens, M. Stegemann, C. R. Glösenkamp, H. D. Volk, C. Goffinet, M. Landthaler, E. Wyler, P. Georg, M. Schneider, C. Dang-Heine, N. Neuwinger, K. Kappert, R. Tauber, V. Corman, J. Raabe, K. M. Kaiser, M. T. Vinh, G. Rieke, C. Meisel, T. Ulas, M. Becker, R. Geffers, M. Witzenrath, C. Drosten, N. Suttorp, C. von Kalle, F. Kurth, K. Händler, J. L. Schultze, A. C. Aschenbrenner, Y. Li, J. Nattermann, B. Sawitzki, A. E. Saliba, L. E. Sander, Severe COVID-19 Is Marked by a Dysregulated Myeloid Cell Compartment. Cell 182, 1419–1440.e1423 (2020).

3. M. L. Meizlish, A. B. Pine, J. D. Bishai, G. Goshua, E. R. Nadelmann, M. Simonov, C. H. Chang, H. Zhang, M. Shallow, P. Bahel, K. Owusu, Y. Yamamoto, T. Arora, D. S. Atri, A. Patel, R. Gbyli, J. Kwan, C. H. Won, C. Dela Cruz, C. Price, J. Koff, B. A. King, H. M. Rinder, F. P. Wilson, J. Hwa, S. Halene, W. Damsky, D. van Dijk, A. I. Lee, H. J. Chun, A neutrophil activation signature predicts critical illness and mortality in COVID-19. Blood Adv 5, 1164–1177 (2021).

4. A. C. Aschenbrenner, M. Mouktaroudi, B. Krämer, M. Oestreich, N. Antonakos, M. Nuesch-Germano, K. Gkizeli, L. Bonaguro, N. Reusch, K. Baßler, M. Saridaki, R. Knoll, T. Pecht, T. S. Kapellos, S. Doulou, C. Kröger, M. Herbert, L. Holsten, A. Horne, I. D. Gemünd, N. Rovina, S. Agrawal, K. Dahm, M. van Uelft, A. Drews, L. Lenkeit, N. Bruse, J. Gerretsen, J. Gierlich, M. Becker, K. Händler, M. Kraut, H. Theis, S. Mengiste, E. De Domenico, J. Schulte-Schrepping, L. Seep, J. Raabe, C. Hoffmeister, M. ToVinh, V. Keitel, G. Rieke, V. Talevi, D. Skowasch, N. A. Aziz, P. Pickkers, F. L. van de Veerdonk, M. G. Netea, J. L. Schultze, M. Kox, M. M. B. Breteler, J. Nattermann, A. Koutsoukou, E. J. Giamarellos-Bourboulis, T. Ulas, Disease severity-specific neutrophil signatures in blood transcriptomes stratify COVID-19 patients. Genome Med 13, 7 (2021).

5. CDC. (https://www.cdc.gov/mis/mis-c/hcp/index.html, 2021), vol. 2021.

6. L. M. Yonker, T. Gilboa, A. F. Ogata, Y. Senussi, R. Lazarovits, B. P. Boribong, Y. C. Bartsch, M. Loiselle, M. N. Rivas, R. A. Porritt, R. Lima, J. P. Davis, E. J. Farkas, M. D. Burns, N. Young, V. S. Mahajan, S. Hajizadeh, X. I. H. Lopez, J. Kreuzer, R. Morris, E. E. Martinez, I. Han, K. Griswold, Jr., N. C. Barry, D. B. Thompson, G. Church, A. G. Edlow, W. Haas, S. Pillai, M. Arditi, G. Alter, D. R. Walt, A. Fasano, Multisystem inflammatory syndrome in children is driven by zonulin-dependent loss of gut mucosal barrier. J Clin Invest 131, (2021).

7. I. Valverde, Y. Singh, J. Sanchez-de-Toledo, P. Theocharis, A. Chikermane, S. Di Filippo, B. Kuciñska, S. Mannarino, A. Tamariz-Martel, F. Gutierrez-Larraya, G. Soda, K. Vandekerckhove, F. Gonzalez-Barlatay, C. J. McMahon, S. Marcora, C. P. Napoleone, P. Duong, G. Tuo, A. Deri, G. Nepali, M. Ilina, P. Ciliberti, O. Miller, Acute Cardiovascular Manifestations in 286 Children With Multisystem Inflammatory Syndrome Associated With COVID-19 Infection in Europe. Circulation 143, 21–32 (2021).

8. L. R. Feldstein, M. W. Tenforde, K. G. Friedman, M. Newhams, E. B. Rose, H. Dapul, V. L. Soma, A. B. Maddux, P. M. Mourani, C. Bowens, M. Maamari, M. W. Hall, B. J. Riggs, J. S. Giuliano, Jr., A. R. Singh, S. Li, M. Kong, J. E. Schuster, G. E. McLaughlin, S. P. Schwartz, T. C. Walker, L. L. Loftis, C. V. Hobbs, N. B. Halasa, S. Doymaz, C. J. Babbitt, J. R. Hume, S. J. Gertz, K. Irby, K. N. Clouser, N. Z. Cvijanovich, T. T. Bradford, L. S. Smith, S. M. Heidemann, S. P. Zackai, K. Wellnitz, R. A. Nofziger, S. M. Horwitz, R. W. Carroll, C. M. Rowan, K. M. Tarquinio, E. H. Mack, J. C. Fitzgerald, B. M. Coates, A. M. Jackson, C. C. Young, M. B. F. Son, M. M. Patel, J. W. Newburger, A. G. Randolph, Characteristics and Outcomes of US Children and Adolescents With Multisystem Inflammatory Syndrome in Children (MIS-C) Compared With Severe Acute COVID-19. Jama 325, 1074–1087 (2021).

9. A. Meghraoui-Kheddar, B. G. Chousterman, N. Guillou, S. M. Barone, S. Granjeaud, H. Vallet, A. Corneau, K. Guessous, C. de Roquetaillade, A. Boissonnas, J. M. Irish, C. Combadière, Two New Neutrophil Subsets Define a Discriminating Sepsis Signature. Am J Respir Crit Care Med, (2021).

10. J. K. Juss, D. House, A. Amour, M. Begg, J. Herre, D. M. Storisteanu, K. Hoenderdos, G. Bradley, M. Lennon, C. Summers, E. M. Hessel, A. Condliffe, E. R. Chilvers, Acute Respiratory Distress Syndrome Neutrophils Have a Distinct Phenotype and Are Resistant to Phosphoinositide 3-Kinase Inhibition. Am J Respir Crit Care Med 194, 961–973 (2016).

11. D. Gómez-Moreno, J. M. Adrover, A. Hidalgo, Neutrophils as effectors of vascular inflammation. Eur J Clin Invest 48 Suppl 2, e12940 (2018).

12. S. Ravichandran, J. Tang, G. Grubbs, Y. Lee, S. Pourhashemi, L. Hussaini, S. A. Lapp, R. C. Jerris, V. Singh, A. Chahroudi, E. J. Anderson, C. A. Rostad, S. Khurana, SARS-CoV-2 immune repertoire in MIS-C and pediatric COVID-19. Nat Immunol 22, 1452–1464 (2021).

13. L. Vella, J. R. Giles, A. E. Baxter, D. A. Oldridge, C. Diorio, L. Kuri-Cervantes, C. Alanio, M. B. Pampena, J. E. Wu, Z. Chen, Y. J. Huang, E. M. Anderson, S. Gouma, K. O. McNerney, J. Chase, C. Burudpakdee, J. H. Lee, S. A. Apostolidis, A. C. Huang, D. Mathew, O. Kuthuru, E. C. Goodwin, M. E. Weirick, M. J. Bolton, C. P. Arevalo, A. Ramos, C. Jasen, H. M. Giannini, D. A. K, N. J. Meyer, E. M. Behrens, H. Bassiri, S. E. Hensley, S. E. Henrickson, D. T. Teachey, M. R. Betts, E. J. Wherry, Deep Immune Profiling of MIS-C demonstrates marked but transient immune activation compared to adult and pediatric COVID-19. medRxiv, (2020).

14. L. E. Hsieh, A. Grifoni, J. Sidney, C. Shimizu, H. Shike, N. Ramchandar, E. Moreno, A. H. Tremoulet, J. C. Burns, A. Franco, Characterization of SARS-CoV-2 and common cold coronavirus-specific T-cell responses in MIS-C and Kawasaki disease children. Eur J Immunol, (2021).

15. A. M. Newman, C. B. Steen, C. L. Liu, A. J. Gentles, A. A. Chaudhuri, F. Scherer, M. S. Khodadoust, M. S. Esfahani, B. A. Luca, D. Steiner, M. Diehn, A. A. Alizadeh, Determining cell type abundance and expression from bulk tissues with digital cytometry. Nat Biotechnol 37, 773–782 (2019).

16. A. Del Prete, L. Martínez-Muñoz, C. Mazzon, L. Toffali, F. Sozio, L. Za, D. Bosisio, L. Gazzurelli, V. Salvi, L. Tiberio, C. Liberati, E. Scanziani, A. Vecchi, C. Laudanna, M. Mellado, A. Mantovani, S. Sozzani, The atypical receptor CCRL2 is required for CXCR2-dependent neutrophil recruitment and tissue damage. Blood 130, 1223–1234 (2017).

17. R. Zilionis, C. Engblom, C. Pfirschke, V. Savova, D. Zemmour, H. D. Saatcioglu, I. Krishnan, G. Maroni, C. V. Meyerovitz, C. M. Kerwin, S. Choi, W. G. Richards, A. De Rienzo, D. G. Tenen, R. Bueno, E. Levantini, M. J. Pittet, A. M. Klein, Single-Cell Transcriptomics of Human and Mouse Lung Cancers Reveals Conserved Myeloid Populations across Individuals and Species. Immunity 50, 1317–1334.e1310 (2019).

18. M. Sundqvist, K. Christenson, A. Holdfeldt, M. Gabl, J. Mårtensson, L. Björkman, R. Dieckmann, C. Dahlgren, H. Forsman, Similarities and differences between the responses induced in human phagocytes through activation of the medium chain fatty acid receptor GPR84 and the short chain fatty acid receptor FFA2R. Biochim Biophys Acta Mol Cell Res 1865, 695–708 (2018).

19. M. Metzemaekers, M. Gouwy, P. Proost, Neutrophil chemoattractant receptors in health and disease: double-edged swords. Cell Mol Immunol 17, 433–450 (2020).

20. M. Reyes, M. R. Filbin, R. P. Bhattacharyya, A. Sonny, A. Mehta, K. Billman, K. R. Kays, M. Pinilla-Vera, M. E. Benson, L. A. Cosimi, D. T. Hung, B. D. Levy, A. C. Villani, M. Sade-Feldman, R. M. Baron, M. B. Goldberg, P. C. Blainey, N. Hacohen, Plasma from patients with bacterial sepsis or severe COVID-19 induces suppressive myeloid cell production from hematopoietic progenitors in vitro. Sci Transl Med 13, (2021).

21. E. J. Gosselin, K. Wardwell, W. F. Rigby, P. M. Guyre, Induction of MHC class II on human polymorphonuclear neutrophils by granulocyte/macrophage colony-stimulating factor, IFN-gamma, and IL-3. J Immunol 151, 1482–1490 (1993).

22. M. J. Carter, M. Fish, A. Jennings, K. J. Doores, P. Wellman, J. Seow, S. Acors, C. Graham, E. Timms, J. Kenny, S. Neil, M. H. Malim, S. M. Tibby, M. Shankar-Hari, Peripheral immunophenotypes in children with multisystem inflammatory syndrome associated with SARS-CoV-2 infection. Nat Med 26, 1701–1707 (2020).

23. J. Fajnzylber, J. Regan, K. Coxen, H. Corry, C. Wong, A. Rosenthal, D. Worrall, F. Giguel, A. Piechocka-Trocha, C. Atyeo, S. Fischinger, A. Chan, K. T. Flaherty, K. Hall, M. Dougan, E. T. Ryan, E. Gillespie, R. Chishti, Y. Li, N. Jilg, D. Hanidziar, R. M. Baron, L. Baden, A. M. Tsibris, K. A. Armstrong, D. R. Kuritzkes, G. Alter, B. D. Walker, X. Yu, J. Z. Li, SARS-CoV-2 viral load is associated with increased disease severity and mortality. Nat Commun 11, 5493 (2020).

24. M. Groselj-Grenc, A. Ihan, M. Derganc, Neutrophil and monocyte CD64 and CD163 expression in critically ill neonates and children with sepsis: comparison of fluorescence intensities and calculated indexes. Mediators Inflamm 2008, 202646 (2008).

25. S. Banerjee, A. Mohammed, H. R. Wong, N. Palaniyar, R. Kamaleswaran, Machine Learning Identifies Complicated Sepsis Course and Subsequent Mortality Based on 20 Genes in Peripheral Blood Immune Cells at 24 H Post-ICU Admission. Front Immunol 12, 592303 (2021).

26. T. E. Andargie, N. Tsuji, F. Seifuddin, M. K. Jang, P. S. Yuen, H. Kong, I. Tunc, K. Singh, A. Charya, K. Wilkins, S. Nathan, A. Cox, M. Pirooznia, R. A. Star, S. Agbor-Enoh, Cell-free DNA maps COVID-19 tissue injury and risk of death and can cause tissue injury. JCI Insight 6, (2021).

27. L. M. Yonker, A. M. Neilan, Y. Bartsch, A. B. Patel, J. Regan, P. Arya, E. Gootkind, G. Park, M. Hardcastle, A. St John, L. Appleman, M. L. Chiu, A. Fialkowski, D. De la Flor, R. Lima, E. A. Bordt, L. J. Yockey, P. D’Avino, S. Fischinger, J. E. Shui, P. H. Lerou, J. V. Bonventre, X. G. Yu, E. T. Ryan, I. V. Bassett, D. Irimia, A. G. Edlow, G. Alter, J. Z. Li, A. Fasano, Pediatric Severe Acute Respiratory Syndrome Coronavirus 2 (SARS-CoV-2): Clinical Presentation, Infectivity, and Immune Responses. J Pediatr 227, 45–52.e45 (2020).

28. H. A. Rothan, S. N. Byrareddy, The potential threat of multisystem inflammatory syndrome in children during the COVID-19 pandemic. Pediatr Allergy Immunol 32, 17–22 (2021).

29. A. K. Mehta, D. T. Gracias, M. Croft, TNF activity and T cells. Cytokine 101, 14–18 (2018).

30. C. de Cevins, M. Luka, N. Smith, S. Meynier, A. Magérus, F. Carbone, V. García-Paredes, L. Barnabei, M. Batignes, A. Boullé, M. C. Stolzenberg, B. P. Pérot, B. Charbit, T. Fali, V. Pirabakaran, B. Sorin, Q. Riller, G. Abdessalem, M. Beretta, L. Grzelak, P. Goncalves, J. P. Di Santo, H. Mouquet, O. Schwartz, M. Zarhrate, M. Parisot, C. Bole-Feysot, C. Masson, N. Cagnard, A. Corneau, C. Brunaud, S. Y. Zhang, J. L. Casanova, B. Bader-Meunier, J. Haroche, I. Melki, M. Lorrot, M. Oualha, F. Moulin, D. Bonnet, Z. Belhadjer, M. Leruez, S. Allali, C. Gras-Leguen, L. de Pontual, A. Fischer, D. Duffy, F. Rieux-Laucat, J. Toubiana, M. M. Ménager, A monocyte/dendritic cell molecular signature of SARS-CoV-2-related multisystem inflammatory syndrome in children with severe myocarditis. Med (N Y) 2, 1072–1092.e1077 (2021).

31. C. Tschöpe, E. Ammirati, B. Bozkurt, A. L. P. Caforio, L. T. Cooper, S. B. Felix, J. M. Hare, B. Heidecker, S. Heymans, N. Hübner, S. Kelle, K. Klingel, H. Maatz, A. S. Parwani, F. Spillmann, R. C. Starling, H. Tsutsui, P. Seferovic, S. Van Linthout, Myocarditis and inflammatory cardiomyopathy: current evidence and future directions. Nat Rev Cardiol 18, 169–193 (2021).

32. I. Müller, T. Vogl, K. Pappritz, K. Miteva, K. Savvatis, D. Rohde, P. Most, D. Lassner, B. Pieske, U. Kühl, S. Van Linthout, C. Tschöpe, Pathogenic Role of the Damage-Associated Molecular Patterns S100A8 and S100A9 in Coxsackievirus B3-Induced Myocarditis. Circ Heart Fail 10, (2017).

33. L. T. Weckbach, U. Grabmaier, A. Uhl, S. Gess, F. Boehm, A. Zehrer, R. Pick, M. Salvermoser, T. Czermak, J. Pircher, N. Sorrelle, M. Migliorini, D. K. Strickland, K. Klingel, V. Brinkmann, U. Abu Abed, U. Eriksson, S. Massberg, S. Brunner, B. Walzog, Midkine drives cardiac inflammation by promoting neutrophil trafficking and NETosis in myocarditis. J Exp Med 216, 350–368 (2019).

34. M. Dolhnikoff, J. Ferreira Ferranti, R. A. de Almeida Monteiro, A. N. Duarte-Neto, M. Soares Gomes-Gouvêa, N. Viu Degaspare, A. Figueiredo Delgado, C. Montanari Fiorita, G. Nunes Leal, R. M. Rodrigues, K. Taverna Chaim, J. R. Rebello Pinho, M. Carneiro-Sampaio, T. Mauad, L. F. Ferraz da Silva, W. Brunow de Carvalho, P. H. N. Saldiva, E. Garcia Caldini, SARS-CoV-2 in cardiac tissue of a child with COVID-19-related multisystem inflammatory syndrome. Lancet Child Adolesc Health 4, 790–794 (2020).

35. A. N. Duarte-Neto, E. G. Caldini, M. S. Gomes-Gouvêa, C. T. Kanamura, R. A. de Almeida Monteiro, J. F. Ferranti, A. M. C. Ventura, F. A. Regalio, D. M. Fiorenzano, M. Gibelli, W. B. Carvalho, G. N. Leal, J. R. R. Pinho, A. F. Delgado, M. Carneiro-Sampaio, T. Mauad, L. F. Ferraz da Silva, P. H. N. Saldiva, M. Dolhnikoff, An autopsy study of the spectrum of severe COVID-19 in children: From SARS to different phenotypes of MIS-C. EClinicalMedicine 35, 100850 (2021).

36. H. Qi, S. Yang, L. Zhang, Neutrophil Extracellular Traps and Endothelial Dysfunction in Atherosclerosis and Thrombosis. Front Immunol 8, 928 (2017).

37. N. L. Denning, M. Aziz, S. D. Gurien, P. Wang, DAMPs and NETs in Sepsis. Front Immunol 10, 2536 (2019).

38. C. M. de Bont, W. C. Boelens, G. J. M. Pruijn, NETosis, complement, and coagulation: a triangular relationship. Cell Mol Immunol 16, 19–27 (2019).

39. L. M. Yonker, J. Boucau, J. Regan, M. C. Choudhary, M. D. Burns, N. Young, E. J. Farkas, J. P. Davis, P. P. Moschovis, T. Bernard Kinane, A. Fasano, A. M. Neilan, J. Z. Li, A. K. Barczak, Virologic Features of Severe Acute Respiratory Syndrome Coronavirus 2 Infection in Children. J Infect Dis 224, 1821–1829 (2021).

40. P. Bastard, A. Gervais, T. Le Voyer, J. Rosain, Q. Philippot, J. Manry, E. Michailidis, H. H. Hoffmann, S. Eto, M. Garcia-Prat, L. Bizien, A. Parra-Martínez, R. Yang, L. Haljasmägi, M. Migaud, K. Särekannu, J. Maslovskaja, N. de Prost, Y. Tandjaoui-Lambiotte, C. E. Luyt, B. Amador-Borrero, A. Gaudet, J. Poissy, P. Morel, P. Richard, F. Cognasse, J. Troya, S. Trouillet-Assant, A. Belot, K. Saker, P. Garçon, J. G. Rivière, J. C. Lagier, S. Gentile, L. B. Rosen, E. Shaw, T. Morio, J. Tanaka, D. Dalmau, P. L. Tharaux, D. Sene, A. Stepanian, B. Megarbane, V. Triantafyllia, A. Fekkar, J. R. Heath, J. L. Franco, J. M. Anaya, J. Solé-Violán, L. Imberti, A. Biondi, P. Bonfanti, R. Castagnoli, O. M. Delmonte, Y. Zhang, A. L. Snow, S. M. Holland, C. Biggs, M. Moncada-Vélez, A. A. Arias, L. Lorenzo, S. Boucherit, B. Coulibaly, D. Anglicheau, A. M. Planas, F. Haerynck, S. Duvlis, R. L. Nussbaum, T. Ozcelik, S. Keles, A. A. Bousfiha, J. El Bakkouri, C. Ramirez-Santana, S. Paul, Q. Pan-Hammarström, L. Hammarström, A. Dupont, A. Kurolap, C. N. Metz, A. Aiuti, G. Casari, V. Lampasona, F. Ciceri, L. A. Barreiros, E. Dominguez-Garrido, M. Vidigal, M. Zatz, D. van de Beek, S. Sahanic, I. Tancevski, Y. Stepanovskyy, O. Boyarchuk, Y. Nukui, M. Tsumura, L. Vidaur, S. G. Tangye, S. Burrel, D. Duffy, L. Quintana-Murci, A. Klocperk, N. Y. Kann, A. Shcherbina, Y. L. Lau, D. Leung, M. Coulongeat, J. Marlet, R. Koning, L. F. Reyes, A. Chauvineau-Grenier, F. Venet, G. Monneret, M. C. Nussenzweig, R. Arrestier, I. Boudhabhay, H. Baris-Feldman, D. Hagin, J. Wauters, I. Meyts, A. H. Dyer, S. P. Kennelly, N. M. Bourke, R. Halwani, N. S. Sharif-Askari, K. Dorgham, J. Sallette, S. M. Sedkaoui, S. AlKhater, R. Rigo-Bonnin, F. Morandeira, L. Roussel, D. C. Vinh, S. R. Ostrowski, A. Condino-Neto, C. Prando, A. Bonradenko, A. N. Spaan, L. Gilardin, J. Fellay, S. Lyonnet, K. Bilguvar, R. P. Lifton, S. Mane, M. S. Anderson, B. Boisson, V. Béziat, S. Y. Zhang, E. Vandreakos, O. Hermine, A. Pujol, P. Peterson, T. H. Mogensen, L. Rowen, J. Mond, S. Debette, X. de Lamballerie, X. Duval, F. Mentré, M. Zins, P. Soler-Palacin, R. Colobran, G. Gorochov, X. Solanich, S. Susen, J. Martinez-Picado, D. Raoult, M. Vasse, P. K. Gregersen, L. Piemonti, C. Rodríguez-Gallego, L. D. Notarangelo, H. C. Su, K. Kisand, S. Okada, A. Puel, E. Jouanguy, C. M. Rice, P. Tiberghien, Q. Zhang, A. Cobat, L. Abel, J. L. Casanova, Autoantibodies neutralizing type I IFNs are present in ~4% of uninfected individuals over 70 years old and account for ~20% of COVID-19 deaths. Sci Immunol 6, (2021).

41. J. L. Jacobs, W. Bain, A. Naqvi, B. Staines, P. M. S. Castanha, H. Yang, V. F. Boltz, S. Barratt-Boyes, E. T. A. Marques, S. L. Mitchell, B. Methé, T. F. Olonisakin, G. Haidar, T. W. Burke, E. Petzold, T. Denny, C. W. Woods, B. J. McVerry, J. S. Lee, S. C. Watkins, C. M. St Croix, A. Morris, M. F. Kearney, M. S. Ladinsky, P. J. Bjorkman, G. Kitsios, J. W. Mellors, SARS-CoV-2 Viremia is Associated with COVID-19 Severity and Predicts Clinical Outcomes. Clin Infect Dis, (2021).

42. Y. Li, A. M. Schneider, A. Mehta, M. Sade-Feldman, K. R. Kays, M. Gentili, N. C. Charland, A. L. Gonye, I. Gushterova, H. K. Khanna, T. J. LaSalle, K. M. Lavin-Parsons, B. M. Lilley, C. L. Lodenstein, K. Manakongtreecheep, J. D. Margolin, B. N. McKaig, B. A. Parry, M. Rojas-Lopez, B. C. Russo, N. Sharma, J. Tantivit, M. F. Thomas, J. Regan, J. P. Flynn, A. C. Villani, N. Hacohen, M. B. Goldberg, M. R. Filbin, J. Z. Li, SARS-CoV-2 viremia is associated with distinct proteomic pathways and predicts COVID-19 outcomes. J Clin Invest 131, (2021).

43. R. Lima, E. F. Gootkind, D. De la Flor, L. J. Yockey, E. A. Bordt, P. D’Avino, S. Ning, K. Heath, K. Harding, J. Zois, G. Park, M. Hardcastle, K. A. Grinke, S. Grimmel, S. P. Davidson, P. J. Forde, K. E. Hall, M. Neilan, J. D. Matute, P. H. Lerou, A. Fasano, J. E. Shui, A. G. Edlow, L. M. Yonker, Establishment of a pediatric COVID-19 biorepository: unique considerations and opportunities for studying the impact of the COVID-19 pandemic on children. BMC Med Res Methodol 20, 228 (2020).

44. A. C. Villani, R. Satija, G. Reynolds, S. Sarkizova, K. Shekhar, J. Fletcher, M. Griesbeck, A. Butler, S. Zheng, S. Lazo, L. Jardine, D. Dixon, E. Stephenson, E. Nilsson, I. Grundberg, D. McDonald, A. Filby, W. Li, P. L. De Jager, O. Rozenblatt-Rosen, A. A. Lane, M. Haniffa, A. Regev, N. Hacohen, Single-cell RNA-seq reveals new types of human blood dendritic cells, monocytes, and progenitors. Science 356, (2017).

45. A. Edwards, W. Haas, Multiplexed Quantitative Proteomics for High-Throughput Comprehensive Proteome Comparisons of Human Cell Lines. Methods Mol Biol 1394, 1–13 (2016).

46. C. S. Hughes, S. Moggridge, T. Müller, P. H. Sorensen, G. B. Morin, J. Krijgsveld, Single-pot, solid-phase-enhanced sample preparation for proteomics experiments. Nat Protoc 14, 68–85 (2019).

47. J. D. Lapek, Jr., P. Greninger, R. Morris, A. Amzallag, I. Pruteanu-Malinici, C. H. Benes, W. Haas, Detection of dysregulated protein-association networks by high-throughput proteomics predicts cancer vulnerabilities. Nat Biotechnol 35, 983–989 (2017).

48. J. Li, J. G. Van Vranken, L. Pontano Vaites, D. K. Schweppe, E. L. Huttlin, C. Etienne, P. Nandhikonda, R. Viner, A. M. Robitaille, A. H. Thompson, K. Kuhn, I. Pike, R. D. Bomgarden, J. C. Rogers, S. P. Gygi, J. Paulo, TMTpro reagents: a set of isobaric labeling mass tags enables simultaneous proteome-wide measurements across 16 samples. Nat Methods 17, 399–404 (2020).

49. G. C. McAlister, D. P. Nusinow, M. P. Jedrychowski, M. Wühr, E. L. Huttlin, B. K. Erickson, R. Rad, W. Haas, S. P. Gygi, MultiNotch MS3 enables accurate, sensitive, and multiplexed detection of differential expression across cancer cell line proteomes. Anal Chem 86, 7150–7158 (2014).

50. L. Ting, R. Rad, S. P. Gygi, W. Haas, MS3 eliminates ratio distortion in isobaric multiplexed quantitative proteomics. Nat Methods 8, 937–940 (2011).

51. B. K. Erickson, M. P. Jedrychowski, G. C. McAlister, R. A. Everley, R. Kunz, S. P. Gygi, Evaluating multiplexed quantitative phosphopeptide analysis on a hybrid quadrupole mass filter/linear ion trap/orbitrap mass spectrometer. Anal Chem 87, 1241–1249 (2015).

52. B. K. Erickson, J. Mintseris, D. K. Schweppe, J. Navarrete-Perea, A. R. Erickson, D. P. Nusinow, J. A. Paulo, S. P. Gygi, Active Instrument Engagement Combined with a Real-Time Database Search for Improved Performance of Sample Multiplexing Workflows. J Proteome Res 18, 1299–1306 (2019).

53. J. K. Eng, A. L. McCormack, J. R. Yates, An approach to correlate tandem mass spectral data of peptides with amino acid sequences in a protein database. J Am Soc Mass Spectrom 5, 976–989 (1994).

54. E. L. Huttlin, M. P. Jedrychowski, J. E. Elias, T. Goswami, R. Rad, S. A. Beausoleil, J. Villén, W. Haas, M. E. Sowa, S. P. Gygi, A tissue-specific atlas of mouse protein phosphorylation and expression. Cell 143, 1174–1189 (2010).

55. J. E. Elias, S. P. Gygi, Target-decoy search strategy for increased confidence in large-scale protein identifications by mass spectrometry. Nat Methods 4, 207–214 (2007).

56. Y. C. Bartsch, C. Wang, T. Zohar, S. Fischinger, C. Atyeo, J. S. Burke, J. Kang, A. G. Edlow, A. Fasano, L. R. Baden, E. J. Nilles, A. E. Woolley, E. W. Karlson, A. R. Hopke, D. Irimia, E. S. Fischer, E. T. Ryan, R. C. Charles, B. D. Julg, D. A. Lauffenburger, L. M. Yonker, G. Alter, Humoral signatures of protective and pathological SARS-CoV-2 infection in children. Nat Med 27, 454–462 (2021).

57. A. Graubert, F. Aguet, A. Ravi, K. G. Ardlie, G. Getz, RNA-SeQC 2: Efficient RNA-seq quality control and quantification for large cohorts. Bioinformatics 37, 3048–3050 (2021).

58. M. I. Love, W. Huber, S. Anders, Moderated estimation of fold change and dispersion for RNA-seq data with DESeq2. Genome Biol 15, 550 (2014).

59. S. V. Puram, I. Tirosh, A. S. Parikh, A. P. Patel, K. Yizhak, S. Gillespie, C. Rodman, C. L. Luo, E. A. Mroz, K. S. Emerick, D. G. Deschler, M. A. Varvares, R. Mylvaganam, O. Rozenblatt-Rosen, J. W. Rocco, W. C. Faquin, D. T. Lin, A. Regev, B. E. Bernstein, Single-Cell Transcriptomic Analysis of Primary and Metastatic Tumor Ecosystems in Head and Neck Cancer. Cell 171, 1611–1624.e1624 (2017).

60. J. Schindelin, I. Arganda-Carreras, E. Frise, V. Kaynig, M. Longair, T. Pietzsch, S. Preibisch, C. Rueden, S. Saalfeld, B. Schmid, J. Y. Tinevez, D. J. White, V. Hartenstein, K. Eliceiri, P. Tomancak, A. Cardona, Fiji: an open-source platform for biological-image analysis. Nat Methods 9, 676–682 (2012).

61. J. Y. Tinevez, N. Perry, J. Schindelin, G. M. Hoopes, G. D. Reynolds, E. Laplantine, S. Y. Bednarek, S. L. Shorte, K. W. Eliceiri, TrackMate: An open and extensible platform for single-particle tracking. Methods 115, 80–90 (2017).

