## Supplemental Figures and Tables for "Neutrophil Profiles of Pediatric COVID-19 and Multisystem Inflammatory Syndrome in Children"

Boribong BP, LaSalle TJ *et al*.

Lael M. Yonker,

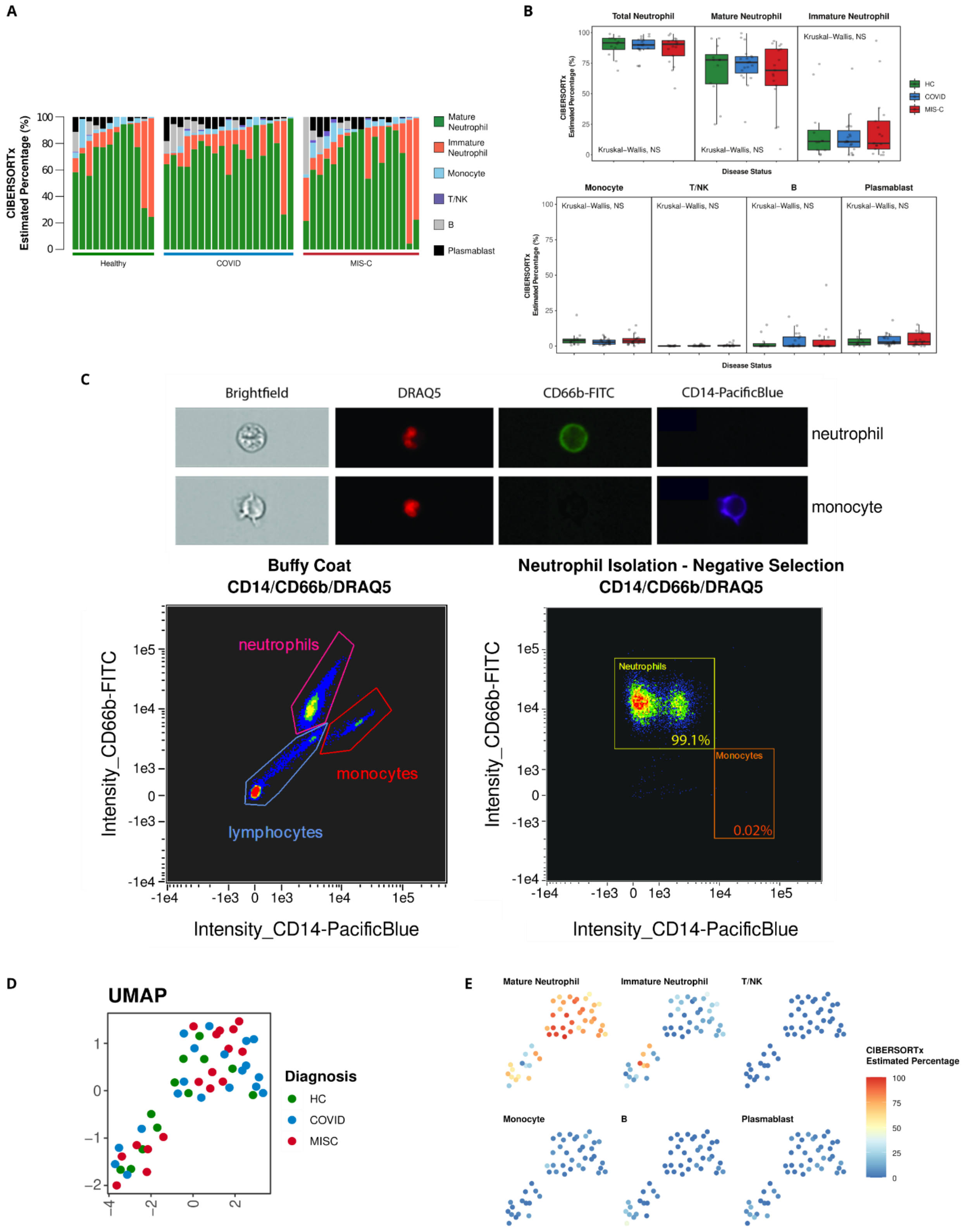
A
CIBERSORTx Estimated Percentage (%)
Mature Neutrophil
Immature Neutrophil
Monocyte
T/NK
B
Plasmablast
Healthy
COVID
MIS-C
B
Total Neutrophil
Mature Neutrophil
Immature Neutrophil
Kruskal–Wallis, NS
HC
COVID
MIS-C
Disease Status
Monocyte
T/NK
B
Plasmablast
C
Brightfield
DRAQ5
CD66b-FITC
CD14-PacificBlue
neutrophil
monocyte
Buffy Coat
CD14/CD66b/DRAQ5
Neutrophil Isolation - Negative Selection
CD14/CD66b/DRAQ5
neutrophils
monocytes
lymphocytes
Neutrophils
99.1%
Monocytes
0.02%
Intensity_CD66b-FITC
Intensity_CD14-PacificBlue
D
UMAP
Diagnosis
HC
COVID
MISC
E
Mature Neutrophil
Immature Neutrophil
T/NK
Monocyte
B
Plasmablast
CIBERSORTx Estimated Percentage

**Fig. S1. Neutrophil isolation and quality control for purity, related to Fig. 1.** (**A**) Bar plots displaying the distribution of the CIBERSORTx estimated cell type percentages for each sample. Samples are divided by disease status and then ordered according to Total Neutrophil Content (sum of Mature and Immature neutrophil fractions). (**B**) Box plots showing the distribution of CIBERSORTx percentages for each cell type divided by disease status. No significant differences were found between disease grouping using the Kruskal-Wallis test. **(C)** Cell are gated on nuclear stain, DRAQ5 (far red). Neutrophils are identified by CD66b (FITC), and monocytes by CD14 (Pacific Blue). Left: population of nucleated cells from the buffy coat. Right: population of isolated neutrophils after negative selection, confirming high purity of neutrophil population. (**D**), (**E**) UMAP plots of neutrophil bulk RNA-seq data. Each point is one bulk sample, and points are color coded according to disease status (**D**) or CIBERSORTx estimated cell type percentages (**E**).

A

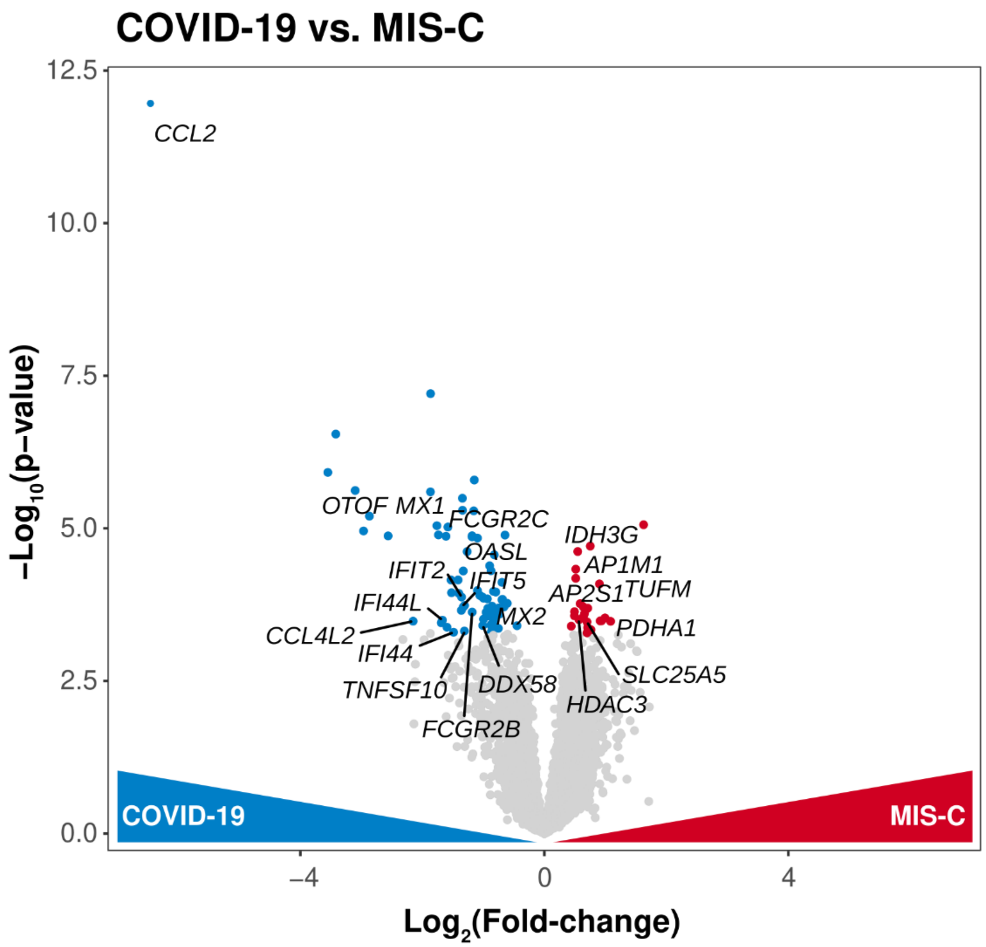

COVID-19 vs. MIS-C
12.5
10.0
7.5
5.0
2.5
0.0
−Log10(p−value)
CCL2
OTOF MX1
FCGR2C
OASL
IFIT2
IFIT5
IFI44L
CCL4L2
IFI44
MX2
TNFSF10
DDX58
FCGR2B
IDH3G
AP1M1
AP2S1
TUFM
PDHA1
SLC25A5
HDAC3
COVID-19
MIS-C
−4
0
4
Log2(Fold-change)


B

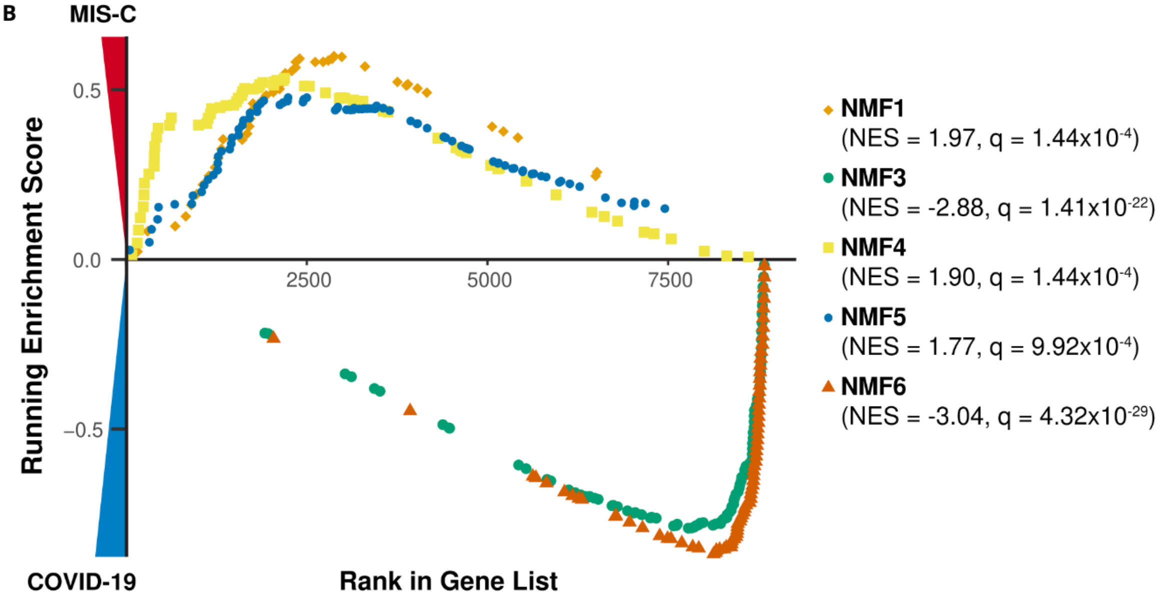

MIS-C
0.5
0.0
−0.5
2500
5000
7500
Running Enrichment Score
COVID-19
Rank in Gene List
NMF1
(NES = 1.97, q = 1.44x10-4)
NMF3
(NES = -2.88, q = 1.41x10-22)
NMF4
(NES = 1.90, q = 1.44x10-4)
NMF5
(NES = 1.77, q = 9.92x10-4)
NMF6
(NES = -3.04, q = 4.32x10-29)

**Fig. S2. Direct comparison of acute pediatric COVID-19 and MIS-C, Related to Fig. 3.** (**A**) Volcano plot showing differentially expressed genes between MIS-C samples and acute pediatric COVID-19 samples. Color-coded points indicate genes that pass FDR correction with $q < 0.05$.(**B**) GSEA enrichment plots for the NMF1 (Pro-Neu), NMF3 (PD-L1+ ISG+), NMF4 (Immature) NMF5 (G-MDSC), and NMF6 (ISG+) signatures, all of which passed FDR correction as indicated.

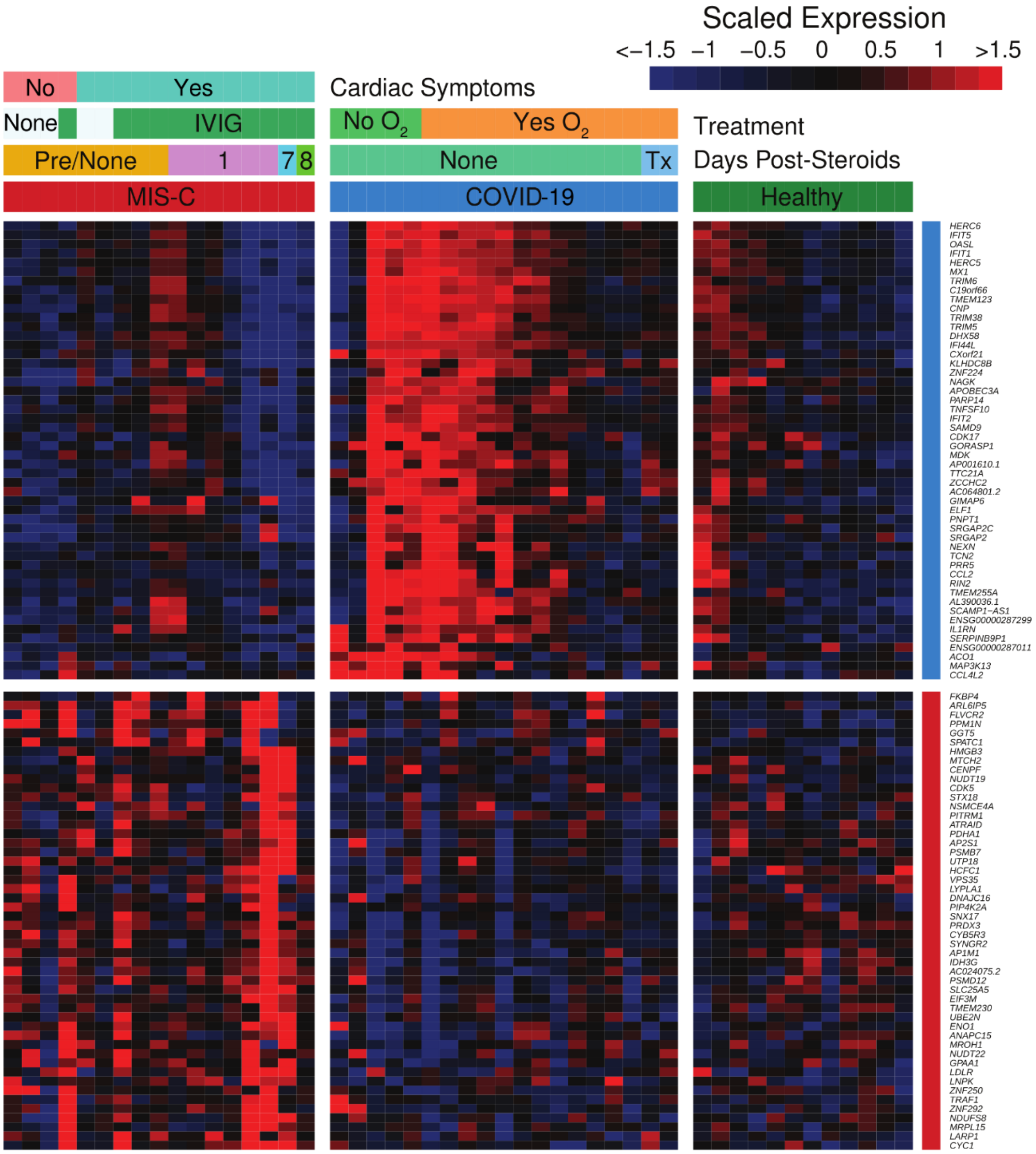


**Fig. S3. Comparison of MIS-C, acute pediatric COVID-19, and healthy children Related to Fig. 3.** Heatmap displaying scaled RNA-seq expression values for gene markers distinguishing pediatric acute COVID-19 and MIS-C. Color-coded bars above indicate disease status, cardiac involvement for MIS-C patients, course of treatment, and days since administration of steroids.

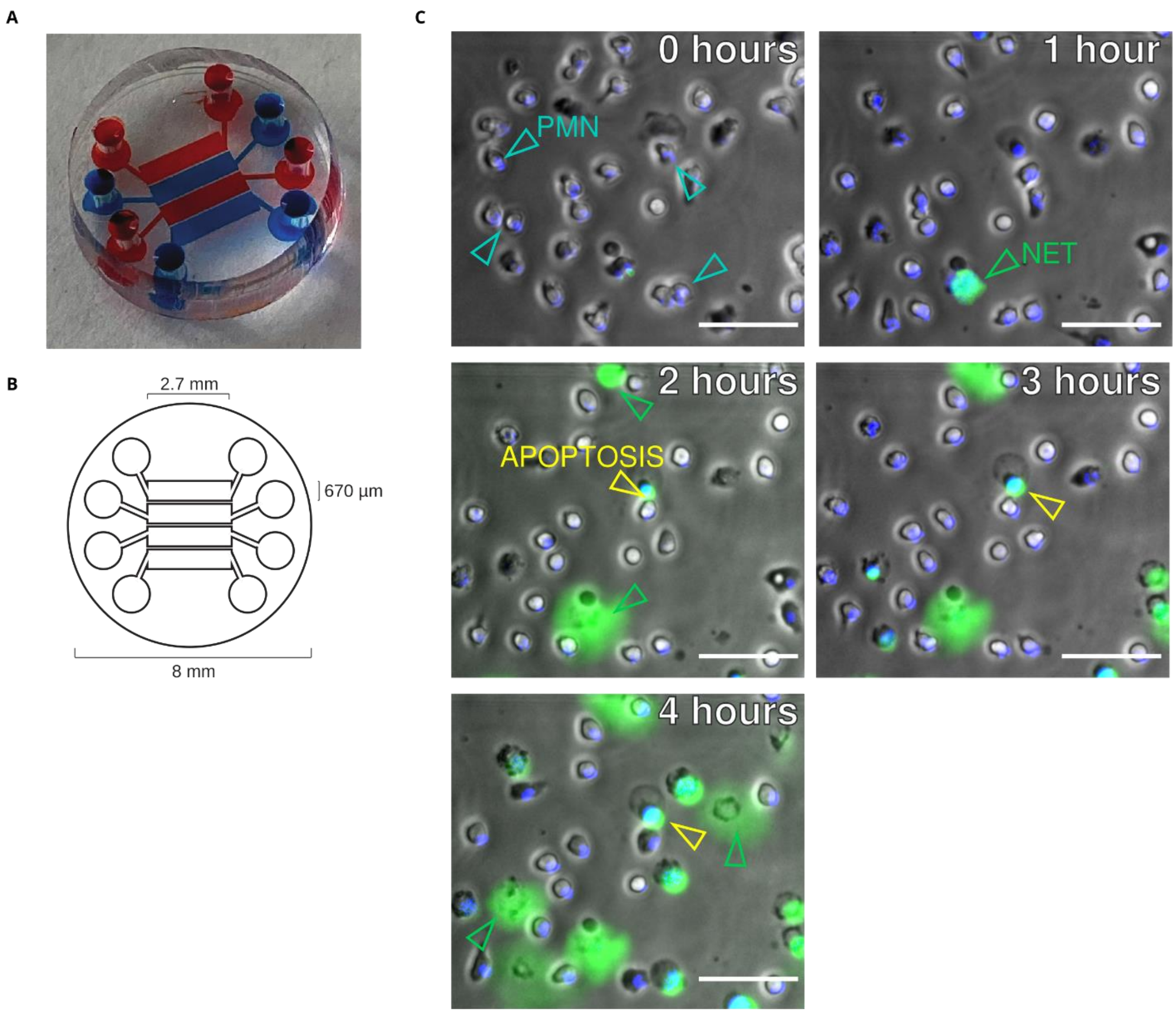


**Fig. S4. Design and schematic of microfluidic device, related to Fig.** 5. (**A**) Microfluidic device used to visualize NET release over time. Dye showing 4 separated channels which allow for 4 simultaneous measurements. (**B**) Dimension of microfluidic device used to visualize NET release over time. (**C**) Time-lapse images of NET formation and apoptosis within the microfluidic devices. Neutrophil (PMN) nucleus was stained with Hoechst stain (DAPI) and extracellular DNA was stained with SYTOX green (FITC). Diffused SYTOX green staining and disrupted membrane seen in the brightfield channel over time was determined to be NETosis. Circular, stable SYTOX green staining and circular neutrophils seen over time were determined to be an apoptotic cell.

Scale bar = 50 µM.

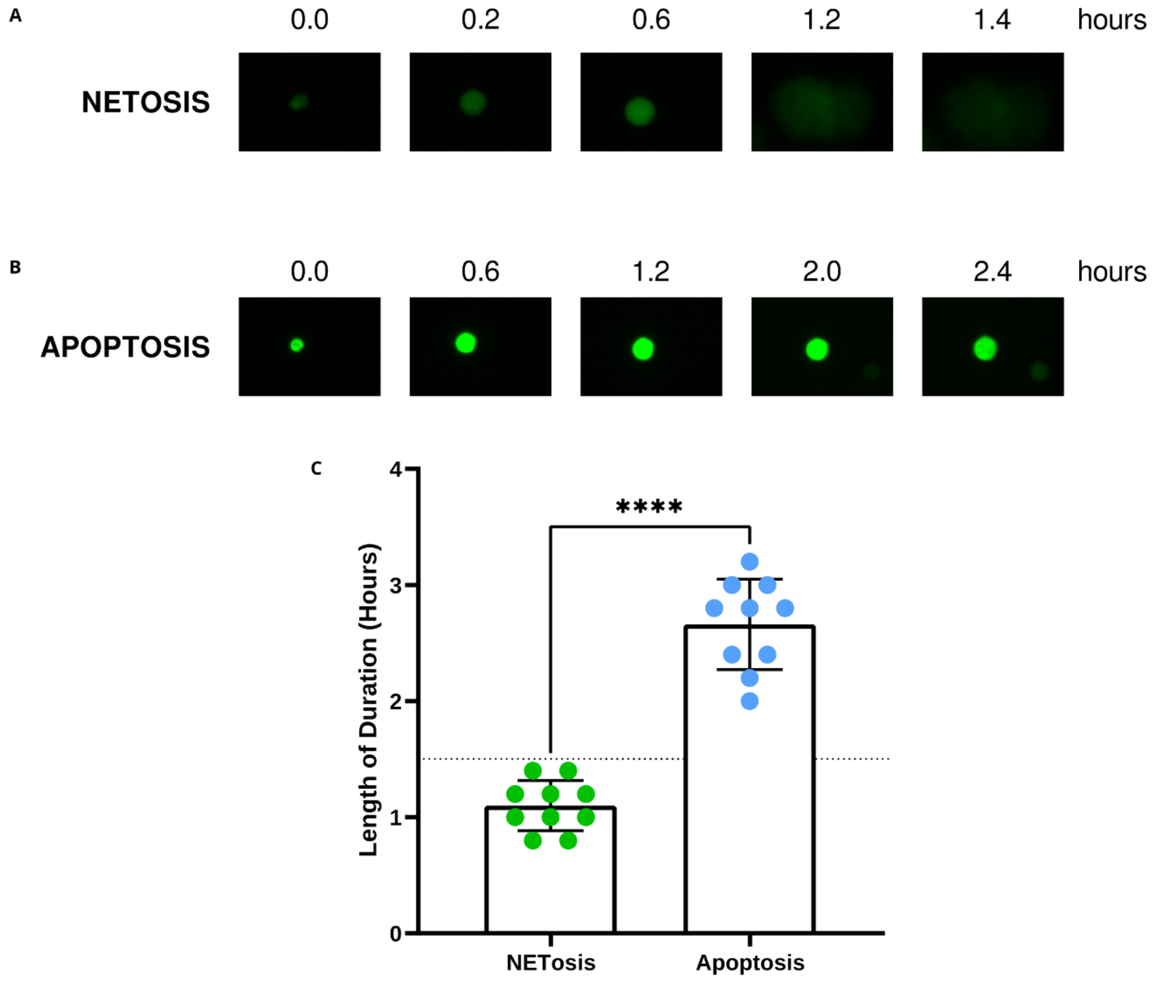


**Fig. S4. Methodology and validation of quantification of NETosis, related to Figs. 4 and 5.** (**A**) Time-lapse visualization of NET release via SYTOX green staining of extracellular DNA (FITC) where the extracellular DNA diffuses over time. (**B**) Time-lapse visualization of apoptosis via SYTOX green staining of extracellular DNA (FITC) where extracellular DNA is circular and stable over time. (**C**) Validation of quantification of NETosis. Neutrophils were manually tracked to determine the length of SYTOX green duration in NET release ($n$ = 10) and apoptosis ($n$ = 10). NETs were determined to be SYTOX green staining that was detected for ≤ 1.5 hrs and apoptotic cells were defined as tracked cells with a duration > 1.5 hrs. Significance was determined by 1-way ANOVA with multiple comparisons in GraphPad Prism v9. Mean values and standard deviation are presented. Statistical significance is defined as **** $P < 0.0001$.

Supplementary Tables

| Patients with acute COVID-19 infection | Age (years) | Sex at Birth | SARS-CoV-2 RT-PCR On Admission | Hospitalized | Highest Level of Care | Treatment Required | Treated with Steroids | Respiratory Support Needed | $O_2$ Needed | Patient Intubated |
|---|---|---|---|---|---|---|---|---|---|---|
| 1 | 10 | M | (+) | yes | ward | no | no | no | no | no |
| 2 | 14 | M | (+) | yes | ward | no | no | no | no | no |
| 3 | 15 | F | (+) | yes | ward | yes | no | yes | yes | no |
| 4 | 0.05 | M | (+) | yes | ward | yes | no | yes | yes | no |
| 5 | 16 | F | (+) | yes | ward | yes | no | yes | yes | no |
| 6 | 2.5 | M | (+) | yes | PICU | yes | no | yes | yes | no |
| 7 | 12.6 | M | (+) | yes | ward | yes | no | yes | yes | yes |
| 8 | 13.81 | M | (+) | yes | PICU | yes | yes | yes | yes | no |
| 9 | 17.15 | M | (+) | yes | ward | no | no | no | no | no |
| 10 | 15.71 | M | (+) | yes | PICU | yes | no | yes | yes | no |
| 11 | 22.16 | M | (+) | yes | ward | yes | no | yes | no | no |
| 12 | 13 | F | (+) | yes | PICU | no | no | no | no | no |
| 13 | 20.35 | M | (+) | yes | ward | yes | yes | no | no | no |
| 14 | 17.59 | F | (+) | yes | PICU | yes | yes | yes | yes | no |
| 15 | 21.99 | F | (+) | yes | ward | yes | yes | yes | yes | no |
| 16 | 21.86 | M | (+) | yes | ward | yes | yes | yes | yes | no |
| 17 | 19.15 | M | (+) | yes | ward | yes | yes | yes | yes | no |
| 18 | 1.55 | M | (+) | yes | ward | yes | no | yes | yes | no |
| 19 | 3.74 | M | (+) | yes | PICU | yes | yes | yes | yes | no |
| 20 | 16.33 | M | (+) | yes | PICU | no | no | yes | yes | no |
| 21 | 0.92 | M | (+) | yes | outpatient | no | no | no | no | no |
| 22 | 18 | F | (+) | no | outpatient | no | no | no | no | no |

| | | | | | | | | | | |
|---|---|---|---|---|---|---|---|---|---|---|
| 23 | 20 | F | (+) | no | outpatient | no | no | no | no | no |
| 24 | 20 | M | (+) | no | outpatient | no | no | no | no | no |
| 25 | 16 | F | (+) | no | outpatient | no | no | no | no | no |
| 26 | 13 | M | (+) | no | outpatient | no | no | no | no | no |
| 27 | 13 | M | (+) | no | outpatient | no | no | no | no | no |
| 28 | 19 | M | (+) | no | outpatient | no | no | no | no | no |
| 29 | 17 | F | (+) | no | outpatient | no | no | no | no | no |
| 30 | 15 | F | (+) | no | outpatient | no | no | no | no | no |
| 31 | 16 | M | (+) | no | outpatient | no | no | no | no | no |
| 32 | 10 | M | (+) | no | outpatient | no | no | no | no | no |
| 33 | 19 | F | (+) | no | outpatient | no | no | no | no | no |
| 34 | 2 | F | (+) | no | outpatient | no | no | no | no | no |
| 35 | 17 | M | (+) | no | outpatient | no | no | no | no | no |
| 36 | 10 | F | (+) | no | outpatient | no | no | no | no | no |
| 37 | 21.4 | M | (+) | no | outpatient | no | no | no | no | no |
| 38 | 11.3 | M | (+) | no | outpatient | no | no | no | no | no |
| 39 | 18 | M | (+) | no | outpatient | no | no | no | no | no |
| 40 | 15.2 | F | (+) | no | outpatient | no | no | no | no | no |
| 41 | 16 | M | (+) | no | outpatient | no | no | no | no | no |
| 42 | 17.3 | M | (+) | no | outpatient | yes | yes | yes | yes | no |
| 43 | 12 | M | (+) | no | outpatient | no | no | no | no | no |

**Table S1.** Demographics and clinical characteristics of pediatric acute COVID-19 patients included in the study.

| Patients with MIS-C | Age (years) | Sex at Birth | SARS-CoV-2 RT-PCR On Admission | SARS-CoV-2 Antibody | COVID-19 Exposure | Highest Level of Care | Cardiovascular Involvement | Treatment Required | Treated with IVIG | Treated with Steroids | Treated with Anikinra |
|---|---|---|---|---|---|---|---|---|---|---|---|
| 1 | 2.9 | M | n/a | n/a | (+) | ward | no | no | no | no | no |
| 2 | 1.2 | M | neg | neg | (+) | ward | no | no | no | no | no |
| 3 | 1 | F | neg | (+) | (+) | ward | no | no | no | no | no |
| 4 | 1.5 | M | neg | (+) | (+) | ward | no | yes | yes | no | no |
| 5 | 17 | F | neg | (+) | (+) | ward | no | yes | no | yes | no |
| 6 | 2 | F | neg | (+) | (+) | ward | no | yes | yes | yes | no |
| 7 | 18 | F | neg | (+) | (+) | ward | no | yes | no | tes | no |
| 8 | 9 | M | neg | neg | (+) | ward | no | no | no | no | no |
| 9 | 0.17 | M | (+) | n/a | (+) | ward | no | no | no | no | no |
| 10 | 0.17 | M | neg | neg | (+) | ward | no | no | no | no | no |
| 11 | 0.17 | F | (+) | n/a | (+) | ward | no | no | no | no | no |
| 12 | 1.34 | M | neg | (+) | (+) | PICU | no | yes | yes | yes | yes |
| 13 | 14.79 | M | neg | (+) | (+) | ward | myocarditis | no | no | no | no |
| 14 | 7.6 | F | n/a | (+) | (+) | PICU | ventricular dysfunction (EF 48%), vasopressor support (epinephrine) | yes | yes | yes | yes |
| 15 | 2.6 | M | neg | neg | (+) | ward | atypical Kawasaki's, lack of coronary tapering | yes | yes | no | no |
| 16 | 13.6 | M | neg | (+) | (+) | PICU | myocarditis | yes | yes | yes | yes |

| | | | | | | | | | | | |
|---|---|---|---|---|---|---|---|---|---|---|---|
| 17 | 12.5 | M | neg | (+) | (+) | PICU | myocarditis, elevated NT-proBNP | yes | yes | yes | yes |
| 18 | 3.5 | M | neg | (+) | (+) | ward | coronary arterial aneurysm | yes | yes | yes | no |
| 19 | 7.11 | M | (+) | n/a | (+) | PICU | coronary aneurysm, elevated troponin, ventricular dyfunction (EF 48) | yes | yes | yes | no |
| 20 | 21 | M | n/a | (+) | (+) | PICU | myocarditis, ventricular dysfunction (EF 21%), vasopressor support | yes | yes | yes | no |
| 21 | 19.4 | F | n/a | (+) | (+) | PICU | extracorporeal membrane oxygenation | yes | yes | yes | no |
| 22 | 10.3 | M | (+) | (+) | (+) | PICU | ventricular dysfunction, vasopressor suppport, Mobitz type I and II | yes | yes | yes | yes |
| 23 | 9.63 | M | n/a | (+) | (+) | PICU | extracorporeal membrane oxygenation | yes | yes | yes | no |
| 24 | 8.27 | M | neg | (+) | (+) | PICU | ventricular dysfunction (EF 40) | yes | yes | yes | no |
| 25 | 5.61 | F | neg | (+) | (+) | ward | ventricular dysfunction (EF 52%); conduction | yes | yes | yes | no |

| | | | | | | | | | | | |
|---|---|---|---|---|---|---|---|---|---|---|---|
| | | | | | | | defect (1st degree AV block) | | | | |
| 26 | 9 | M | neg | (+) | (+) | ward | elevated BNP, coronary dilation or aneurysm | yes | yes | yes | yes |
| 27 | 3 | F | (+) | (+) | (+) | PICU | bradycardia | yes | no | yes | no |
| 28 | 9 | M | neg | (+) | (+) | PICU | elevated troponin and pro-BNP | yes | yes | no | no |
| 29 | 7 | M | neg | (+) | (+) | PICU | dilated left ventricle | yes | yes | yes | yes |
| 30 | 4 | M | (+) | (+) | (+) | ward | corornary dilation or aneurysm | yes | yes | yes | no |
| 31 | 8 | M | neg | (+) | (+) | PICU | hypotension, ventricular failure, extracorporeal membrane oxygenation, arrhythmia, elevated troponin, elevated bnp, coronary dilation or aneurysm | yes | yes | yes | yes |

**Table S2.** Demographics and clinical characteristics of pediatric MIS-C patients included in the study.

**Movie S1.** Spontaneous NETosis of neutrophils isolated from a healthy pediatric patient captured by fluorescence microscopy (brightfield and FITC). Extracellular DNA stained with SYTOX green. Scale bar = 100 μM.

**Movie S2.** Spontaneous NETosis of neutrophils isolated from a child with MIS-C captured by fluorescence microscopy (brightfield and FITC). Extracellular DNA stained with SYTOX green. Scale bar = 100 μM.

**Movie S3.** NETosis of neutrophils isolated from a healthy pediatric patient stimulated with 100 nM PMA captured by fluorescence microscopy (brightfield and FITC). Extracellular DNA stained with SYTOX green. Scale bar = 100 μM.

**Movie S4.** NETosis of isolated neutrophils isolated from a healthy patient stimulated with PBS-treated Spike beads captured by fluorescence microscopy (brightfield and FITC). Extracellular DNA stained with SYTOX green. Scale bar = 100 μM.

**Movie S5.** NETosis of isolated neutrophils isolated from a healthy patient stimulated with non-COVID-19 plasma-treated Spike beads captured by fluorescence microscopy (brightfield and FITC). Extracellular DNA stained with SYTOX green. Scale bar = 100 μM.

**Movie S6.** NETosis of isolated neutrophils isolated from a healthy patient stimulated with convalescent COVID-19 plasma captured by fluorescence microscopy (brightfield and FITC). Extracellular DNA stained with SYTOX green. Scale bar = 100 μM.

**Movie S7.** NETosis of isolated neutrophils isolated from a healthy patient stimulated with convalescent COVID-19 plasma-treated Spike beads captured by fluorescence microscopy (brightfield and FITC). Extracellular DNA stained with SYTOX green. Scale bar = 100 μM.

**Movie S8.** NETosis of isolated neutrophils isolated from a healthy patient stimulated with MIS-C plasma captured by fluorescence microscopy (brightfield and FITC). Extracellular DNA stained with SYTOX green. Scale bar = 100 μM.

**Movie S9.** NETosis of isolated neutrophils isolated from a healthy patient stimulated with MIS-C plasma-treated Spike beads captured by fluorescence microscopy (brightfield and FITC). Extracellular DNA stained with SYTOX green. Scale bar = 100 μM.
